# Non-neuronal, TGF-β–driven extracellular matrix restructuring promotes neurodegeneration in a PSP-Richardson syndrome model

**DOI:** 10.1101/2025.08.17.670724

**Authors:** Clara Zannino, Desirèe Valente, Davide Bressan, Andreas Bruzelius, Caterina Gabriele, Stefania Scalise, Raffaele Covello, Giorgia Lucia Benedetto, Mariagrazia Talarico, Vittorio Abbonante, Andrea Quattrone, Marco Gaspari, Aldo Quattrone, Fulvio Chiacchiera, Alessandra Fiorenzano, Giovanni Cuda, Elvira Immacolata Parrotta

## Abstract

Progressive supranuclear palsy–Richardson syndrome (PSP-RS) is a rapidly progressive tauopathy lacking effective therapies. Although tau aggregation is a defining feature, the initiating mechanisms remain elusive. Here we used patient-derived induced pluripotent stem cell midbrain organoids, integrating single-cell transcriptomics, bulk RNA profiling, and quantitative proteomics, to dissect early pathogenic events. We identified vascular leptomeningeal-like cells (VLMCs) as the first altered population, exhibiting TGF-β–driven extracellular matrix (ECM) remodeling enriched in collagens, integrins, and TGFBI. The resulting pathological ECM increased stiffness, induced integrin clustering, and activated RhoA-ROCK-mediated cytoskeletal disorganization. These changes sustained PI3K-AKT and MAPK-ERK signaling, suppressed PP2A, hyperactivated mTOR, and impaired autophagy, culminating in tau hyperphosphorylation and mislocalization. Pharmacological inhibition of TGF-β, AKT, ERK, or mTORC1 restored autophagic flux, reduced tau burden, and rescued neuronal architecture. Our findings establish non-neuronal, matrix-producing niche cells as upstream drivers of tauopathy and reveal TGF-β–mediated ECM restructuring as a mechanochemical trigger of neurodegeneration, opening multiple therapeutic avenues for PSP-RS and related tauopathies.

## Background

Progressive supranuclear palsy-Richardson syndrome (PSP-RS) is the most common clinical variant of progressive supranuclear palsy (PSP), a primary 4-repeat (4R) tauopathy ^1^ and a leading cause of atypical parkinsonism. PSP-RS typically manifest in the sixth decade of life and is characterized by early postural instability, vertical gaze palsy, axial rigidity, and cognitive decline, progressing relentlessly and lacking effective treatments ^2,3,4^. Neuropathologically, it is characterized by the widespread deposition of hyperphosphorylated tau in neurons and glia, including tufted astrocytes as a defining lesion ^5,6^. Although tau dysregulation has been extensively studied in Alzheimer’s disease (AD) and other tauopathies ^7^, the molecular events that initiate tau aggregation in sporadic PSP-RS remain poorly defined^,9^, representing a critical barrier to disease-modifying interventions ^10,11^. Historically, tauopathy research has centered on neuron-intrinsic stress pathways. Yet the brain is an integrated multicellular ecosystem, and pathological processes in non-neuronal compartments can exert decisive influence over neuronal fate. In peripheral tissues, vascular niche cells regulate homeostasis and repair through extracellular matrix (ECM) remodeling ^12^ and growth factor signaling ^13^, mechanisms that, if dysregulate, can drive chronic degeneration ^14^. Whether analogous vascular-matrix pathways operate in the central nervous system, and whether they contribute to tauopathy initiation, has remained unexplored. Recent single-nucleus RNA sequencing studies, such as Whitney et al. ^15^, have analyzed post-mortem PSP brain tissue to reveal cell-type specific transcriptomic signatures, highlighting dysregulation of the integrated stress response and EIF2 signaling in neurons, astrocytes, and oligodendrocytes, and their correlation with tau pathology. In contrast, we investigated PSP-RS pathogenesis using patient-derived midbrain organoids and interated multi-omics approaches (single-cell RNA sequencing, bulk transcriptomics, and quantitative proteomics), allowing us to capture early, potentially causal molecular events preceding overt neurodegeneration. We identified a previously unrecognized role for vascular-leptomeningeal-like cells (VLMCs), a perivascular stromal population, in initiating neurodegeneration through aberrant TGF-β signaling that drives ECM remodeling ^16,17^. Strikingly, we previously demonstrated that TGF-β1 is markedly elevated in neuron-derived extracellular vesicles from PSP-RS patients, with minimal or no overlap with other parkinsonian disorders or healthy controls ^18^. This signaling cascade remodels ECM, alters tissue mechanics, and activates neuronal integrin-FAK-kinase pathways, culminating in phosphatase suppression, autophagic failure, and tau accumulation and mislocalization. By reframing PSP-RS pathogenesis around an extracellularly orchestrated cascade, our findings establish a paradigm in which vascular niche cells act as active initiators of tauopathy rather than passive bystanders. This perspective highlights the vascular-matrix interface as a therapeutic entry point for PSP-RS and potentially other neurodegenerative disorders.

## Methods

### Generation of induced pluripotent stem cells (iPSCs)

All experiments were performed in strict accordance with institution, national, and international ethical guidelines and regulations. This study was conducted in full accordance with applicable ethical guidelines and regulations. Human peripheral blood samples were obtained from four individuals with sporadic PSP-RS and three age- and sex-matched healthy controls after written informed consent and approval of the *“Comitato Etico Territoriale Regione Calabria”* (approval #143, 13 May 2024) and the ethics committee of the Azienda Ospedaliero-Universitaria “Renato Dulbecco”. iPSCs were reprogrammed from peripheral blood mononuclear cells (PBMCs) using the CytoTune™-iPS 2.0 Sendai Reprogramming Kit (Thermo Fisher Scientific, Waltham, MA, USA), following the manufacturer’s instruction. The derivation and validation of PSP-RS and control iPSC lines, including assessment of pluripotency and trilineage differentiation potential, have been previously described ^19^. The HC_001 line was established independently and characterized separately ^20^. All lines were confirmed to be free of Sendai virus, mycoplasma, and karyotypically stable prior to use in downstream differentiation and omics experiments.

### Generation of midbrain organoids (MOs)

Midbrain organoids (MOs) were generated from iPSCs derived from four individuals with sporadic PSP–Richardson syndrome and three healthy controls, following an established differentiation protocol ^19,21^. iPSCs were dissociated using Accutase and pooled at a 1:1 ratio (10,000 cells per aggregate) prior to seeding in ultra-low attachment 96-well U-bottom plates (Corning, NY, USA) in mTeSR1 Plus medium supplemented with 10 µM Y-27632 (Miltenyi Biotec, Bergisch Gladbach, Germany). Embryoid bodies (EBs) were differentiated toward a midbrain fate in N2 base medium containing 10 µM SB431542, 100 ng/mL Noggin, 300 ng/mL SHH-C24II, and 1.5 µM CHIR99021 (all from Miltenyi Biotec) from day 0 to day 7. On day 8, medium was replaced with N2 supplemented with 100 ng/mL FGF-8b. From day 11, organoids were maintained in B27 base medium supplemented with 100 ng/mL FGF-8b, 20 ng/mL BDNF (Miltenyi Biotec), and 200 µM L-ascorbic acid (Sigma-Aldrich). On day 14, organoids were embedded in Matrigel (Corning, NY, USA) and cultured in B27 medium containing 20 ng/mL BDNF, 10 ng/mL GDNF (R&D Systems, Bio-Techne, Minneapolis, MN, USA), 200 µM L-ascorbic acid, 500 µM cAMP (Sigma-Aldrich), and 1 µM DAPT (Tocris, Bio-Techne, Minneapolis, MN, USA) to promote dopaminergic maturation. To ensure consistency, organoids were derived from at least three independent differentiations per condition, with a minimum of 50 organoids per batch used for downstream analyses.

### Generation of ventral midbrain dopaminergic neurons (vDAs)

Ventral midbrain dopaminergic (vDA) neurons were generated from iPSCs as described by Nolbrant et al. ^22^, with minor modifications during the early patterning. iPSCs from healthy controls and PSP-RS patients were dissociated using Accutase, pooled at a 1:1 ratio, and seeded at a 2 × 10⁴ cells per well in Laminin-111–coated 24-well plates (1 μg/cm²; Biolamina, Sundbyberg, Sweden) in N2 base medium supplemented with 10 μM Y-27632, 10 μM SB431542, 100 ng/mL Noggin, 300 ng/mL SHH-C24II, and 0.6 μM CHIR99021 (Miltenyi Biotec). Y-27632 was withdrawn on day 2. From day 9, cells were cultured in N2 medium supplemented with 100 ng/mL FGF8. On day 11, cultures were dissociated and replated at 8 × 10⁵ cells/cm² onto Laminin-111–coated wells in B27 medium containing 10 μM Y-27632, 100 ng/mL FGF8, 20 ng/mL BDNF (Miltenyi Biotec), and 0.2 mM L-ascorbic acid (Sigma-Aldrich). Y-27632 was removed on day 14. Neural identity was verified between days 14–16. For terminal differentiation, vDA progenitors were dissociated on day 16 and seeded at 2 × 10⁵ cells/cm² onto wells coated with 2 μg/cm² Laminin-111. Maturation was carried out in B27 medium containing 20 ng/mL BDNF, 10 ng/mL GDNF (R&D Systems), 0.2 mM L-ascorbic acid, 500 μM db-cAMP (Sigma-Aldrich), and 1 μM DAPT (Tocris).

### Whole-cell patch-clamp electrophysiology

Whole-cell patch-clamp recordings were obtained from DA progenitors between day 45-50 of differentiation. Coverslips with cells were transferred to a recording chamber continually perfused with artificial cerebrospinal fluid (aCSF) containing (in mM): 119 NaCl, 2.5 KCl, 1.3 MgSO_4_, 2.5 CaCl_2_, 1.25 NaH_2_PO_4_, 25 Glucose and 26 NaHCO_3_, and gassed with 95% O2–5% CO2 (carbogen gas) at a RT (pH ∼7.4, osm 305 ∼mOsm). Patch pipettes (5–7 MΩ resistance) were pulled from standard borosilicate glass by using a micropipette puller. Recording pipettes were filled with intracellular solution containing (in mM): 122.5 K-gluconate, 12.5 KCl, 0.2 EGTA, 10 HEPES, 2 MgATP, 0.3 Na_3_GTP, and 8 NaCl adjusted to pH 7.3 with KOH and 290–295 mOsm. Whole-cell patch-clamp recordings were obtained using a Multiclamp 700B amplifier and pClamp 10.4 data acquisition software. Access resistance was monitored through the recording, to make sure it did not increase significantly. Resting membrane potential (RMP) was measured, immediately after breaking into the cell. The capacitance of the cell was directly obtained from pClamp 10.4 data acquisition software. Evoked action potentials (AP) were recorded from cells kept around −60 mV through current injection, stepwise current was applied from −20 to 60 pA in at 20 pA increments to push cells towards threshold and elicit induced Aps. The AP threshold, peak amplitude, after hyperpolarisation and AP halfwidth were measured from the first observed spike reaching a minimum of 10 mV and with a duration less than 10 ms evoked by the rheobase current injection step. Sodium and potassium currents were evoked by stepwise injection of depolarizing potentials from −70 mV to +40 mV (10 mV increments). Spontaneous synaptic activity was recorded in voltage-clamp mode at −70 mV. Data were analysed using Clampfit 10.3 (Axon Instruments, Molecular Devices, USA), Mini Analysis Program (Synaptosoft) and Igor Pro 8.04 (Wavemetrics, Portland, OA, USA) combined with the NeuroMatic package ^23^.

### RNA extraction and quantitative real-time PCR

Total RNA was isolated using TRIzol reagent (Thermo Fisher Scientific) according to the manufacturer’s protocol. cDNA was synthesized from 1 μg total RNA using the High-Capacity cDNA Reverse Transcription Kit (Thermo Fisher Scientific). Quantitative PCR was performed using SensiFAST SYBR Hi-ROX chemistry (Meridian Bioscience) on a QuantStudio™ 7 Pro system (Applied Biosystems). Gene expression was normalized to *GAPDH* as an endogenous reference. Relative transcript levels were calculated using the ΔΔCt method. Primer sequences are listed in Supplementary Table 1.

### Immunofluorescence staining and imaging

Organoids were fixed in 4% paraformaldehyde (PFA) for 5 h at room temperature, incubated overnight at 4 °C in 30% (w/v) sucrose (Santa Cruz Biotechnology), and equilibrated in a 1:1 mixture of OCT embedding medium and 30% sucrose for 6 hours at 4 °C. Samples were embedded in OCT (Avantor) and cryosectioned at 20 μm using a cryostat. Sections were post-fixed in 4% PFA for 10 min, permeabilized, and blocked in PBS with 5% goat serum (Thermo Fisher Scientific) and 0.3% Triton X-100 (Sigma-Aldrich) for 1 h at room temperature. Primary antibodies were applied overnight at 4 °C in blocking buffer, followed by incubation with Alexa Fluor 488- or 594-conjugated secondary antibodies (Thermo Fisher Scientific) for 1 h at room temperature in the dark. Nuclei were counterstained with DAPI (Thermo Fisher Scientific), and sections were mounted in Fluoromount aqueous medium (Sigma-Aldrich). Imaging was performed on a Leica Mica microscope. Multiple regions of interest per sample were acquired. Quantification of fluorescence intensity, puncta density, and spatial mapping of neuroanatomical features was performed using Fiji/ImageJ, including the Neuroanatomy Shortcuts plugin. Antibody details are listed in Supplementary Table 2.

### Protein extraction and immunoblotting

Organoids were lysed in ice-cold RIPA buffer (150 mM NaCl, 1% Triton X-100, 0.5% sodium deoxycholate, 0.1% SDS, 50 mM Tris-HCl, pH 7.5) supplemented with Halt™ protease and phosphatase inhibitors (Thermo Fisher Scientific). Lysates were sonicated (10 cycles, 30 s ON/30 s OFF; Diagenode Bioruptor) and incubated on ice for 30 min. Supernatants were collected after centrifugation at 21,000 × g for 1 hour at 4 °C. Protein concentration was determined using the Bradford assay (Bio-Rad). Equal amounts of protein (20 μg per lane) were denatured at 70 °C for 10 min in LDS sample buffer with reducing agent (Thermo Fisher Scientific), separated on 4–12% Bis-Tris Plus SDS-PAGE gels (Thermo Fisher Scientific), and transferred to nitrocellulose membranes (Bio-Rad) using the Trans-Blot Turbo system. Membranes were blocked for 1 h at room temperature in TBS-T with 5% non-fat dry milk (PanReac AppliChem), incubated overnight at 4 °C with primary antibodies, and probed with HRP-conjugated secondary antibodies (Jackson ImmunoResearch) for 1 h at room temperature. Detection was performed using Clarity™ ECL substrate (Bio-Rad) and visualized with the Alliance™ Q9-Atom system (Uvitec). Band intensities were quantified using ImageJ. GAPDH or histone H3 served as loading controls. Antibodies are listed in Supplementary Table 3. Original, uncropped images are included in the Source Data.

### Quantification and statistical analysis

Statistical analyses were performed using GraphPad Prism (v9.3.1). Normality of datasets was assessed using Shapiro–Wilk, D’Agostino–Pearson, Anderson–Darling, or Kolmogorov–Smirnov tests. For comparisons between two groups, unpaired two-tailed t-tests with Welch’s correction were applied to normally distributed data; otherwise, the non-parametric Mann–Whitney U test was used. Statistical tests are specified in the figure legends. Data are presented as mean ± standard error of the mean (s.e.m.) and reflect at least two or three independent biological replicates. All experiments included technical replicates and a minimum of three organoids per condition from at least three independent iPSC differentiation batches. Western blot quantifications were based on n = 3 biological replicates. Immunofluorescence analyses (n > 3) included regional quantification of mean fluorescence intensity, puncta density, and neuroanatomical parameters. Synaptic density was quantified as the number of yellow puncta per cell, reflecting colocalization of presynaptic and PSD95-positive postsynaptic markers, using a validated ImageJ-based colocalization method ^24^. Neuronal morphology was assessed by quantifying neurite branch number and average branch length. Statistical significance was defined as follows: *P* < 0.05 (\**), P < 0.01 (**), P **<** 0.001 (**\*\*\****), *P* < 0.0001 (****).

### Bulk RNA sequencing and analysis

#### RNA extraction and library preparation

Total RNA was extracted from day-90 midbrain organoids derived from pooled iPSCs of three healthy controls and four PSP-RS patients (three biological replicates per group). Poly(A)+ mRNA was purified using oligo(dT) magnetic beads, fragmented, and reverse-transcribed using random hexamer primers. Strand-specific libraries were generated by incorporating dUTP during second-strand synthesis, as described previously ^25^, followed by end repair, adaptor ligation, size selection, USER enzyme digestion, PCR amplification, and purification. Non-strand-specific libraries followed the same protocol without the USER digestion step. Libraries were quantified via Qubit fluorometry and real-time PCR, and fragment size was assessed using a Bioanalyzer (Agilent Technologies). Sequencing was performed on an Illumina platform, with libraries pooled according to effective molarity and target read depth.

#### Preprocessing and alignment

Raw reads (n = 6; three PSP-RS and three HC samples) were quality-trimmed using *fastp* v0.20.0 ^26^. High-quality reads were aligned to the human GRCh38 reference genome using STAR v2.7.3 ^27^, with annotations from GENCODE v42. PCR duplicates were removed using samblaster with the --removeDups option ^28^. Gene-level counts were obtained using featureCounts v1.6.4 ^29^, using the parameters -t exon -g gene_name.

#### Differential expression and enrichment analysis

Statistical analysis was performed using DESeq2 v1.32 ^30^. Shrinkage of log₂ fold changes were applied using the apeglm method ^31^, and *P*-values were adjusted for multiple testing using independent hypothesis weighting (IHW) ^32^. Differentially expressed genes (DEGs) were defined as those with *adjusted P* < 0.05 and absolute log₂ fold change > 1.5. In total, 430 genes were significantly upregulated and 518 downregulated in PSP-RS relative to controls. Principal component analysis (PCA) confirmed clear segregation of biological replicates by condition. Functional enrichment was performed using clusterProfiler v4.12.2 ^33^. Over-representation analysis (ORA) was conducted on the DEG list, and Gene Set Enrichment Analysis (GSEA) was performed using the Hallmark gene sets (H collection) from the Molecular Signatures Database (MSigDB).

### Quantitative proteomics by mass spectrometry

#### Protein extraction and enzymatic digestion

Total protein was extracted from four independent biological replicates per condition using the RIPA buffer, as previously described. For digestion, 20 µg of protein per sample was diluted in RIPA buffer (final volume: 32 µL). Proteins were reduced with 3.2 µL of 100 mM DTT (1 h, 37 °C), alkylated with 3.8 µL of 200 mM iodoacetamide (1 h, 37 °C), and quenched with 0.6 µL of 100 mM DTT (1 h, 37 °C). Half of each sample (10 µg) was processed using the Protein Aggregation Capture (PAC) method ^34^. Briefly, 10 µg of MagReSyn Hydroxyl magnetic beads (5 µL) were washed twice in 70% acetonitrile (ACN) and incubated for 10 min at 1100 rpm. After sequential washes in ACN and 70% ethanol, protein-bound beads were resuspended in 50 µL of 50 mM triethylammonium bicarbonate (TEAB) and digested overnight at 37 °C (1100 rpm) with Trypsin/Lys-C (200 ng; 1:50 enzyme-to-substrate ratio). Residual peptides were recovered with 50 µL of 0.1% formic acid (FA). Unless otherwise stated, reagents were from Sigma-Aldrich.

#### Liquid chromatography–mass spectrometry (LC–MS/MS)

Peptides were analyzed on an Orbitrap Exploris 480 mass spectrometer (Thermo Fisher Scientific) coupled to an EASY-nLC 1200 system. Peptides were loaded onto a 17 cm capillary column (75 µm i.d.) packed in-house with 3 µm C18 beads (Dr. Maisch). Peptide separation was performed using a 140-minute binary gradient at 300 nL/min: 3–25% buffer B over 90 min, 25–40% over 30 min, 40–100% over 8 min, followed by a 10-min hold at 100% B (buffer A: 2% ACN, 0.1% FA; buffer B: 80% ACN, 0.1% FA). Nanoelectrospray ionization was achieved at 2,000 V.

#### Data acquisition and analysis

Data-independent acquisition (DIA) was used with 30 variable-width isolation windows covering m/z 350–1010, including 23 windows of 15 m/z, 4 of 30 m/z, and 3 of 50 m/z with 0.5 m/z overlap. Full MS scans were acquired at 60,000 resolution; DIA scans at 30,000 resolution (AGC target: 5e5; injection time: 50 ms; normalized collision energy: 25). Raw DIA data were analyzed using Spectronaut v18.7 against a composite database comprising the UniProt human reference proteome (79,740 entries, October 2022) and a Matrigel-derived mouse database (4,074 entries, UniProt Mus musculus, December 2022), yielding a combined search space of 83,814 entries. A 46-protein contaminant database was also included. Protein intensities were log₂-transformed and filtered to retain proteins detected in ≥3 samples per condition. Imputation was used for missing values.

Differentially expressed proteins were identified using a two-sample *t*-test with permutation-based FDR (FDR = 0.05, S₀ = 0.2) in Perseus v2.0.11.0. Proteins of Matrigel origin were excluded post hoc. A total of 369 proteins were significantly regulated (q < 0.05, |log₂FC| > 0.5), of which 135 were upregulated and 221 downregulated in PSP-RS relative to healthy controls.

#### Pathway enrichment analysis

Gene Ontology (GO) and KEGG pathway enrichment analyses were conducted using clusterProfiler v4.12.2 ^35^. Gene Set Enrichment Analysis (GSEA) was performed against KEGG gene sets to identify pathways significantly enriched in PSP-RS organoids.

### Drug treatments and pathway modulation

All pharmacological interventions were performed on MOs or vDA neurons derived from at least three independent iPSC differentiation batches per condition. Treatments began at day 90 of MOs maturation (day 100 for vDA neurons) and were maintained for two weeks, except for chloroquine and Torin1, which were applied for 24 hours (**Supplementary Data S1c**). To activate TGFβ signaling, control-derived MOs were exposed to a cocktail of TGFβ-1, TGFβ-2, and TGFβ-3 isoforms (ProteinTech, Chicago, IL, USA) at a final concentration of 20 ng/mL. In PSP-RS organoids, TGFβ pathway inhibition was achieved using the selective ALK5 inhibitor SB431542 (10 µM; Miltenyi Biotec). ROCK signaling was blocked using Y-27632 (10 µM; Miltenyi Biotec). Additional downstream pathways were targeted as follows: LY294002 (100 µM; Merck, Darmstadt, Germany) was used to inhibit PI3K/AKT signaling, PD98059 (25 µM; Selleckem, Houston, TX, USA) was employed to block MAPK/ERK signaling, and Torin-1 (1 µM; Selleckem) was applied to inhibit mTOR signaling. For autophagy blockade, organoids were treated with chloroquine (1 mM; Sigma-Aldrich) for 24 hours. In monolayer cultures of PSP-RS-derived vDA neurons, pharmacological treatments began at day 100 of differentiation and continued for two weeks under the same alternate-day regimen. Final concentrations in vDA cultures were: SB431542 (10 µM), LY294002 (20 µM), and Torin-1 (100 nM). All compounds were prepared and administered in accordance with manufacturers’ protocols. A full schematic of the experimental treatment timeline is provided in Extended Data File 1c.

### Whole-genome sequencing and variant analysis

Genomic DNA was extracted from donor samples and sheared into short fragments. End-repair, A-tailing, and ligation to full-length Illumina adapters were performed prior to size selection. Unless otherwise specified, libraries were PCR-amplified and purified using the AMPure XP system (Beckman Coulter, Beverly, MA, USA). Final libraries were evaluated on a Fragment Analyzer (Agilent Technologies), and concentrations were determined using both Qubit fluorometry (Thermo Fisher Scientific) and qPCR. Qualified libraries were pooled and sequenced on Illumina platforms according to effective library concentration and desired sequencing depth. Raw reads were filtered to remove: (i) reads with adapter contamination >10 bp (≤10% mismatches allowed), (ii) reads containing >10% undefined bases (N), and (iii) reads with >50% of bases below Phred quality score 5. High-quality reads were aligned to the human reference genome (hg38) using BWA ^36^. Variant calling was performed using the HaplotypeCaller module of the Genome Analysis Toolkit (GATK) ^37^. Germline SNPs were filtered using GATK’s VariantFiltration tool with the following criteria: quality by depth (QD) < 2.0, Fisher strand bias (FS) > 60.0, mapping quality (MQ) < 40.0, HaplotypeScore > 13.0, MappingQualityRankSum < –12.5, or ReadPosRankSum < –8.0. Variants not meeting these criteria were excluded from further analysis. VCF files were normalized using bcftools norm, splitting multiallelic sites relative to the hg38 reference genome. Filtering was performed using bcftools view with the flags -f., PASS -a -c 1: nonmajor to retain high-confidence, polymorphic variants, followed by additional filtering with bcftools filter to include only variants with allele frequency >0.05 and QD >10, in accordance with demuxlet recommendations. Final variant call files were sorted, indexed, and intersected with RefSeq exon coordinates (hg38) using bedtools intersect to retain only exonic variants for downstream analysis.

### Single-cell RNA sequencing and analysis

Single-cell suspensions were prepared in phosphate-buffered saline containing 0.04% bovine serum albumin, filtered through a 40 μm cell strainer (Biologix), and counted using a LUNA-II™ automated counter (Logos Biosystems). Cell capture and barcoding were performed using the Chromium Single Cell G Chip Kit and Chromium Controller (10x Genomics), following the manufacturer’s protocol. Libraries were prepared using the Chromium Single Cell 3′ Library & Gel Bead Kit v3.1 (10x Genomics). cDNA quality was assessed using High Sensitivity D5000 ScreenTape on the Agilent 4200 TapeStation, and final libraries were quality-checked using High Sensitivity DNA D1000 ScreenTape (Agilent Technologies) before sequencing on the NovaSeq 6000 platform (Illumina). FASTQ files were generated from raw BCL data using bclconvert. Reads were aligned to the GRCh38/hg38 human reference genome and quantified using Cell Ranger v8.0.1 (10x Genomics) with default settings. Ambient RNA contamination was removed using SoupX v1.6.2. Cells were retained if they expressed between 300 and 5000 genes and contained <15% mitochondrial RNA content. Genetic demultiplexing was performed using demuxlet (v1) ^38^, based on exonic SNPs, with alpha values between 0 and 0.5. Doublets (probability >0.5) were excluded. Single-cell libraries were integrated using the IntegrateLayers function in Seurat (v5.1.0), applying the Harmony method for batch correction ^39^. Two donors (HC1 and PSP2) were excluded due to poor integration with other samples. Dimensionality reduction was performed using PCA on the top 30 principal components. Cell clusters were identified and annotated via FindTransferAnchors and TransferData functions in Seurat, using a published human midbrain reference dataset ^40^, and refined with canonical markers ^41^. Pseudo-bulk expression matrices were generated using AggregateExpression in Seurat, treating each donor as a biological replicate. Differential expression analysis was carried out using FindMarkers. Functional enrichment analysis was conducted on differentially expressed genes using the clusterProfiler package (v4.12.2) ^33,35^.

## Results

### Single-cell analysis reveals VLMC expansion and ECM-driven reprogramming in PSP-RS

To model early molecular events in PSP-RS, we generated midbrain organoids (MOs) from induced pluripotent stem cells (iPSCs) of four patients with sporadic PSP-RS and three age- and sex-matched healthy controls (HCs). Whole-genome sequencing excluded known pathogenic variants ^42^. To minimize donor variability, lines were pooled at the embryoid body (EB) stage (**Supplementary Data S1a**). All lines differentiated efficiently into midbrain-like tissues, exhibiting robust expression of regional markers (**Supplementary Fig. S1a-g**) and generated ventral dopaminergic (vDA) neurons (**Supplementary Data S1b; Supplementary Fig. S1h-k).** Whole-cell patch-clamp recordings showed that both HC- and PSP-RS–derived vDA neurons fired spontaneously at rest and generated action potentials upon depolarizing current injection, consistent with a mature dopaminergic phenotype (**Supplementary Fig. S2a-b**). Passive and active membrane properties, including resting potential, AP threshold and amplitude, afterhyperpolarization, capacitance, ionic currents, AP half-width and firing frequency, were comparable between groups (**Supplementary Fig. S2c-g**). PSP-RS organoids recapitulated key pathological features of the disease, including loss of dopaminergic neurons, 4R-tau accumulation, neurofibrillary tangles, and gliosis^22^ validating the model system. To resolve early, cell type-specific alterations in PSP-RS, we performed single-cell RNA sequencing (scRNA-seq) on day-90 midbrain organoids derived from patients with sporadic PSP-RS and HCs. After stringent quality control, demultiplexing, and exclusion of doublets and ambient RNA contamination, we obtained 11,343 high-quality single-cell profiles with comparable metrics across conditions (**Supplementary Fig. S3a-c**). Unsupervised clustering and UMAP dimensionality reduction identified six major populations: neurons, astrocytes, oligodendrocyte precursor cells (OPCs), vascular leptomeningeal-like cells (VLMCs), floor plate progenitors (FPPs), and cycling progenitors (**Fig. 1a-b; Supplementary Fig. S3d**). PSP-RS MOs showed a striking expansion of VLMCs with a concomitant reduction of neuronal and progenitor populations (**Fig. 1c**). VLMCs were enriched for collagen and matrix-related transcripts, including *COL1A1*, *FBLN1*, and *LUM* (**Fig. 1d**), consistent with a role in ECM remodeling. Gene ontology (GO) enrichment revealed repression of neurogenesis and synaptic programs alongside induction of kinase, adhesion, and ECM organization pathways (**Fig. 1e-f**). Within VLMCs, we observed marked upregulation of integrin ligands and focal adhesion genes (**Fig. 1g**).

**Fig. 1.**
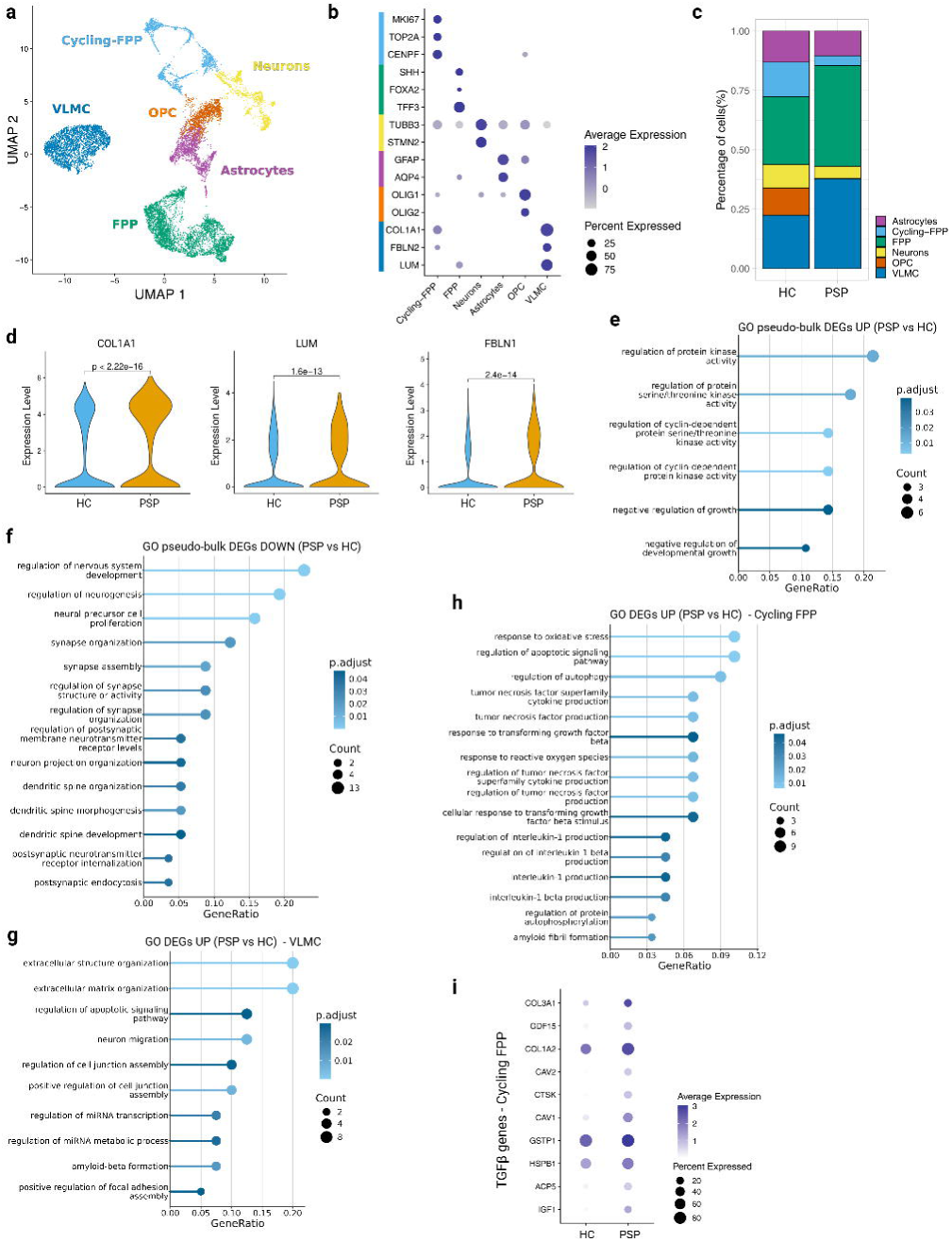
VLMC-driven TGF-β–mediated ECM remodeling merges as the earliest molecular signature of PSP-RS. **a,** UMAP projection of single-cell transcriptomes from HC and PSP-RS MOs identifies seven transcriptionally distinct populations, including neurons, astrocytes, oligodendrocyte precursor cells (OPCs), vascular leptomeningeal cells (VLMCs), fetal progenitor populations (FPP), and cycling-FPPs. **b,** Dot plot showing cell-type–specific expression of canonical marker genes across major clusters. **c,** Stacked bar plot indicates altered cell-type proportions in PSP-RS, with increased representation of VLMCs and cycling-FPPs. **d,** Violin plots show elevated expression of ECM-related genes (*COL1A1, LUM, FBLN1*) in PSP-RS organoids (Wilcoxon test, P < 2.2 × 10⁻¹⁶). **e, f,** GO enrichment analysis of pseudo-bulk differentially expressed genes (DEGs) in PSP-RS versus HC reveals upregulation of pathways involved in cell growth and protein kinase activity (**e**) and downregulation of neurogenesis, synapse organization, and neuronal differentiation (**f**). **g,** GO terms enriched in DEGs upregulated in PSP-RS VLMCs include ECM organization, junction assembly, focal adhesion, and amyloid-β formation. **h,** DEGs in cycling-FPP cells show enrichment for oxidative stress responses, autophagy, cytokine signaling, and TGF-β–responsive pathways. **i,** Dot plot showing increased expression of TGF-β target genes, including *COL3A1*, *COL6A2*, *CSPG4*, and *IGF1*, in cycling-FPP cells from PSP-RS organoids relative to HC.

Transcriptomic alterations extended to other non-neuronal compartments. Cycling FPPs upregulated oxidative stress– and autophagy-related genes, many of which are TGF-β–responsive (*COL3A1*, *CAV1/2, IGFBP3*, *GDF15*, *GSTM1*) (**Fig. 1h-i, Supplementary Fig. S3e**), while astrocytes and progenitors increased expression of *COL1A2* and *FBLN1,* respectively (**Supplementary Fig. S3f**). Cell-type-specific analyses revealed activation of ER stress, unfolded protein response (UPR), and apoptotic programs in FPPs ^43^ (**Supplementary Fig. S3g**), chaperone and IRE1-mediated UPR programs in astrocytes ^44^ (**Supplementary Fig. S3h**), and actin cytoskeleton reorganization and UPR genes in neurons ^45,46^ (**Supplementary Fig. S3i**). These data establish VLMC expansion and ECM-associated transcriptional reprogramming as an early hallmark of PSP-RS, positioning vascular niche cells as upstream coordinated of multicellular stress responses.

### VLMC-driven TGF-β signaling and ECM restructuring as an early pathogenic event

To assess whether VLMC expansion was associated with structural matrix changes, we performed bulk transcriptomic and quantitative proteomic profiling at day 90. PSP-RS MOs exhibited strong molecular divergence from HC (**Supplementary Fig. S4a-b**), with 430 genes upregulated and 518 downregulated in PSP-RS (adjusted *p* < 0.05; **Fig. 2a**), and 356 differentially expressed proteins (135 upregulated, 221 downregulated; **Fig. 2b**). GO analysis identified consistent enrichment for ECM organization, TGF-β–related signaling, and kinase activation (**Fig. 2c-d**). Among the most upregulated proteins was TGFBI, a TGF-β–inducible ECM component (**Fig. 2e**). Heatmap confirmed coordinated upregulation of fibrous collagens and matrix regulators (COL1A1/2, COL3A1, COL6A1, FBLN1, LOX2, SMOC1) (**Fig. 2f-g**). Immunoblotting validated increased levels of COL6A1 and COL1A1 proteins in PSP-RS MOs (**Supplementary Fig. S4c**). Gene Set Enrichment Analysis (GSEA) revealed significant upregulation of ECM organization (NES = 1.935; **Fig. 2h**), collagen formation (NES = 1.927, *p* = 8.47 × 10⁻⁵), focal adhesion (NES = 1.587, *p* = 0.0053), and ECM–receptor interaction (NES = 1.512, *p* = 0.015; **Supplementary Fig. S4d-f**). Consistent with integrin engagement, PSP-RS MOs exhibited increased vinculin-positive focal adhesion puncta (**Fig. 2i**). These findings identify TGF-β–driven ECM remodeling as an early pathogenic signature downstream of VLMC expansion, establishing the structural framework for subsequent intracellular signaling cascades.

**Fig. 2.**
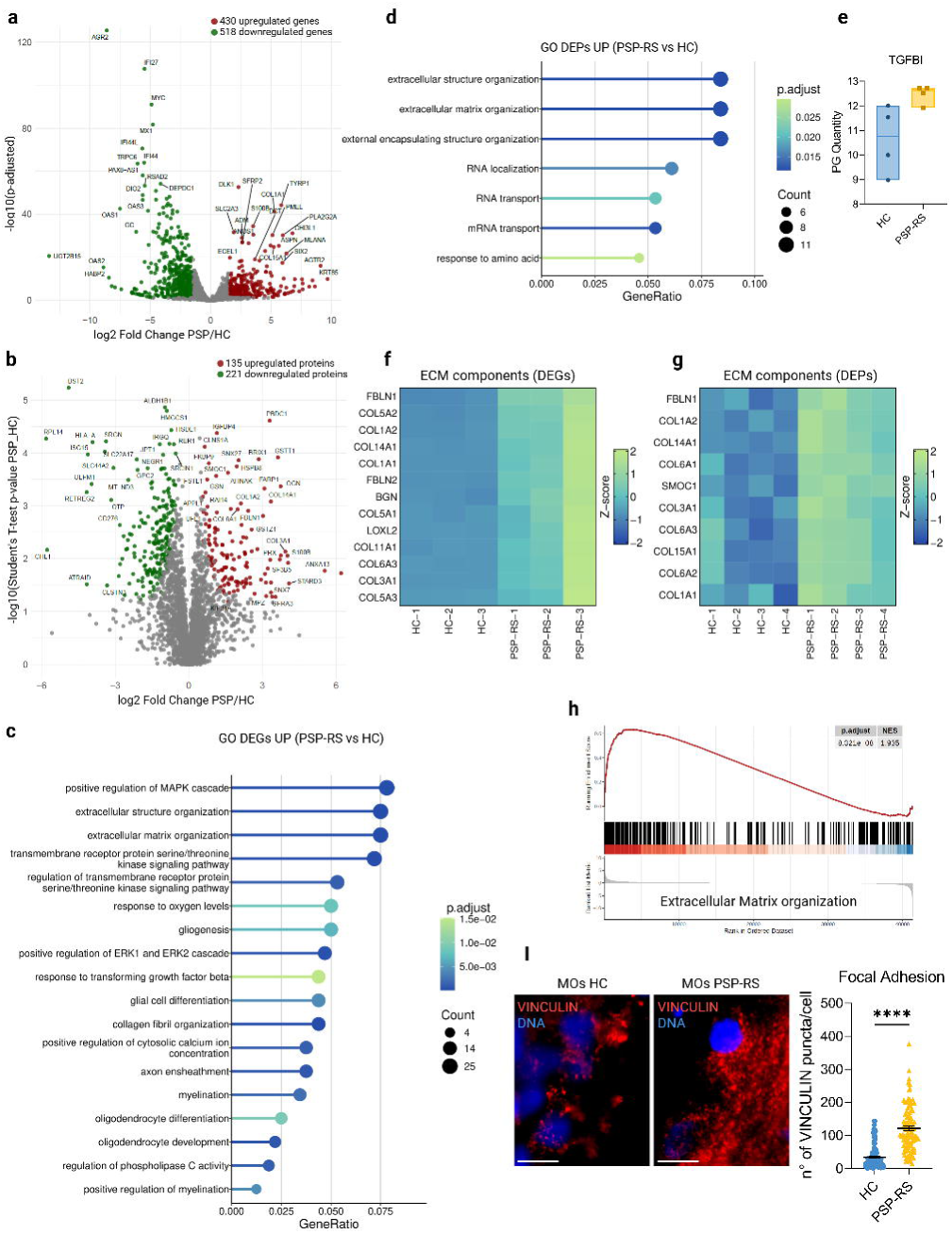
Multi-omic profiling identify ECM remodeling and focal adhesion as defining features of PSP-RS. **a,** Volcano plot of differentially expressed genes (DEGs) in PSP-RS (n = 4) versus healthy control (HC, n = 3) midbrain organoids (MOs); significantly upregulated (red) and downregulated (green) transcripts are indicated (adjusted P < 0.05). **b,** Volcano plot of differentially expressed proteins (DEPs) from proteomic profiling of PSP-RS (n = 4) versus HC (n = 3); significantly upregulated (red) and downregulated (green) proteins are shown (Student t-test P value < 0.05). **c,** Gene ontology (GO) enrichment analysis of DEGs upregulated in PSP-RS MOs reveals enrichment for extracellular matrix (ECM) organization, MAPK cascade activation, and oligodendrocyte-related pathways. **d,** GO enrichment analysis of upregulated DEPs identifies similar enrichment in extracellular structure organization and RNA-associated processes. **e,** Quantification of TGFBI protein abundance (PG quantity) from proteomic datasets shows increased levels in PSP-RS MOs relative to HC. **f, g,** Heatmaps showing Z-score–normalized expression of selected ECM-related components at the transcript (**f**) and protein (**g**) levels across individual samples. **h,** Gene set enrichment analysis (GSEA) of transcriptomic data shows positive enrichment of the “extracellular matrix organization” gene set in PSP-RS MOs (normalized enrichment score [NES] = 1.595; false discovery rate [FDR] q = 0.01). **i,** Representative immunofluorescence images of primary MO-derived cells stained for the focal adhesion marker VINCULIN (red) and nuclei (DAPI, blue) from HC and PSP-RS lines. Scale bars, 10 μm. Quantification of VINCULIN-positive puncta per cell demonstrates a significant reduction in PSP-RS (**** P < 0.0001, Mann–Whitney test; mean ± s.e.m.; n = 101 cells). Statistics are available in Additional File 2.

### Pathological ECM stiffness engages integrin-AKT-ERK pathways to drive tau pathology

Pathway analysis of transcriptomic and proteomic datasets revealed enrichment of PI3K/AKT and MAPK/ERK pathways (**Supplementary Fig. S5a**), consistent with integrin-mediated activation during non-canonical known TGF-β signaling ^47,48^. Immunoblotting confirmed increased phosphorylation of AKT (Ser473) and ERK1/2 (Thr202/Tyr204) (**Fig. 3a-b; Supplementary Fig. S5b-c)**, coupled with reduced expression of PP2A (**Fig. 3c; Supplementary Fig. S5d**), a phosphatase that negatively regulates both pathways and dephosphorylates tau ^49,50^. Pharmacological inhibition of PI3K using LY294002 reduced AKT activation and decreased tau phosphorylation at Ser396 and Thr181 (**Fig. 3d**), while MEK inhibition with PD0325901 suppressed ERK activity and tau phosphorylation (**Fig. 3e**). These pathways operated independently (**Supplementary Fig. S5e-f**). Inhibition of TGF-β receptor I with SB431542 suppressed phosphorylation of both kinases and reduced tau phosphorylation at Ser396 and T181 (**Fig. 3f; Supplementary Fig. S5g**), placing them downstream of TGF-β signaling ^51,52^. Thus, ECM remodeling emerges as the upstream driver of integrin-kinase signaling cascades converging on tau hyperphosphorylation.

**Fig. 3.**
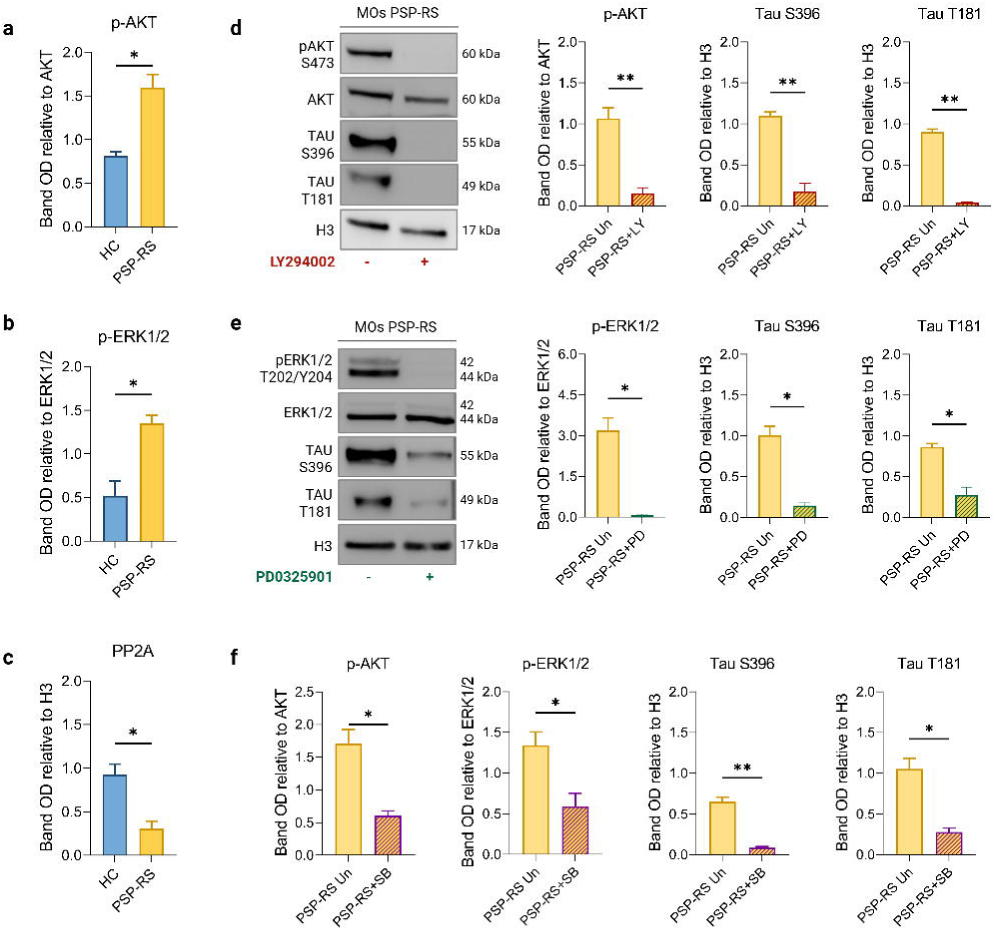
ECM–integrin signaling activates parallel AKT and ERK cascades converging on tau phosphorylation. **a, b,** Immunoblot quantification of phosphorylated AKT (p-AKT^S473) and ERK1/2 (p-ERK1/2^T202/Y204) in MOs from HC and PSP-RS donors shows increased activation of both pathways in PSP-RS (mean ± s.e.m.; n = 3; P = 0.0269 for p-AKT, P = 0.0203 for p-ERK1/2; unpaired t-test with Welch’s correction). **c,** Expression of the PP2A phosphatase is significantly reduced in PSP-RS MOs (P = 0.0179). **d,** Treatment of PSP-RS MOs with the PI3K inhibitor LY294002 reduces p-AKT levels and diminishes phosphorylation of tau at S396 and T181 (P = 0.0081, P = 0.0051, and P = 0.0014, respectively). **e,** Inhibition of ERK1/2 signaling with PD0325901 leads to decreased p-ERK1/2 and reduced phospho-tau at S396 and T181 in PSP-RS MOs (P = 0.0204, P = 0.0103, and P = 0.0146, respectively). **f,** Pharmacological blockade of TGF-β signaling with SB431542 reduces p-AKT and p-ERK1/2, as well as phospho-tau at S396 and T181 in PSP-RS cultures (P = 0.0255, P = 0.0314, P = 0.0059, and P = 0.0167, respectively). Band intensities are normalized to total protein or histone H3 (H3) as loading controls. Uncropped western blot gels and statistics are available in Additional File 1-2.

### mTORC1 hyperactivation impairs autophagy, promoting tau accumulation and mislocalization

Given AKT activation, we examined mTORC1 signaling, a key inhibitor of autophagy ^53^. PSP-RS MOs exhibited elevated phosphorylation of mTOR (Ser2448) and its canonical effectors (p70S6K, RPS6, 4EBP1) (**Figure 4a; Supplementary Fig. S6a**), confirming sustained mTORC1 activation. This was accompanied by p62/SQSTM1 accumulation (**Figure 4b-d; Supplementary Fig. S6b**) and LC3 flux blockade under chloroquine treatment (**Supplementary Fig. S6c-d**). Inhibition of lysosomal degradation in HC MOs phenocopied tau hyperphosphorylation (**Supplementary Fig. S6e-f)**, confirming that impaired proteostasis is sufficient to drive tau accumulation in healthy controls. Pharmacological inhibition of mTORC1 with Torin-1 reduced phosphorylation of p70S6K and 4EBP1 (**Supplementary Fig. S6g-h**) and significantly decreased tau phosphorylation in PSP-RS MOs (**Fig. 4e**; **Supplementary Fig. S6i**), mirroring the effects of upstream AKT inhibition. Notably, tau mislocalized into MAP2-positive compartments in PSP-RS vDA neurons (**Fig. 4f-g**), consistent with pathological redistribution during proteostasis suppression ^54,55,56^. Treatment with LY294002 (PI3K inhibitor) or Torin-1 restored somatodendritic tau compartmentalization and reduced dendritic tau burden (**Fig. 4h-i**). These findings define proteostatic failure as a downstream consequence of VLMC-TGF-β–ECM axis signaling.

**Fig. 4.**
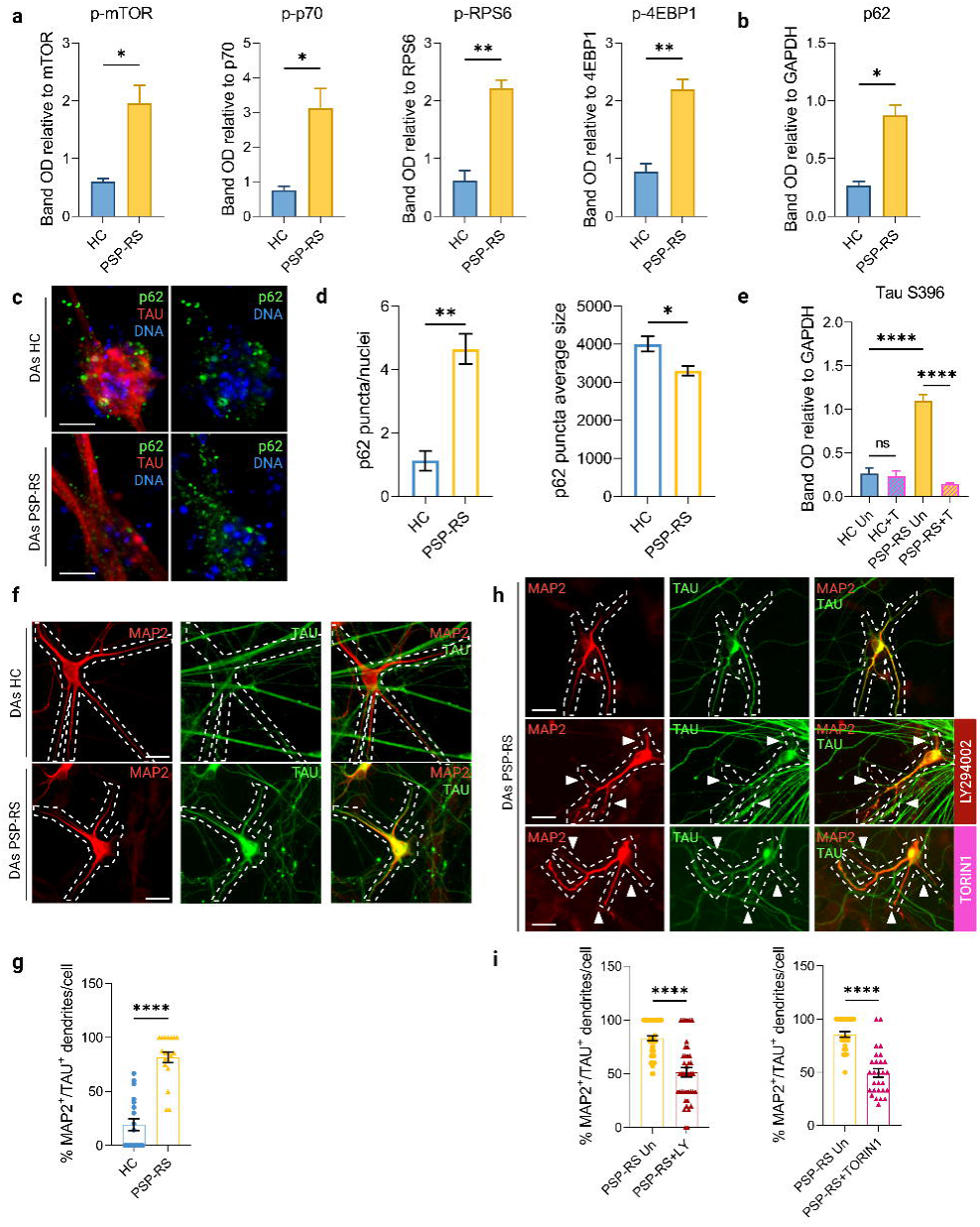
Autophagy impairment through AKT–mTORC1 hyperactivation fuels tau accumulation and mislocalization. **a,** Immunoblot analysis of MOs lysates from HC and PSP-RS donors reveals increased phosphorylation of mTOR and its downstream targets p70^S6K, RPS6, and 4EBP1 in PSP-RS, consistent with mTORC1 hyperactivation (mean ± s.e.m.; n = 3; P < 0.05 for p-mTOR and p-p70^S6K, P < 0.01 for p-RPS6 and p-4EBP1; unpaired t-test with Welch’s correction). **b,** Western blot quantification shows elevated levels of the autophagy adaptor p62 in PSP-RS MOs (P = 0.0116), indicating impaired autophagic flux. **c, d,** Representative confocal images and quantification of p62 and TAU in differentiated dopaminergic (DA) neurons show increased number and size of p62-positive puncta in PSP-RS compared to HC (mean ± s.e.m.; n = 4 images; P = 0.0014 and P = 0.0306, respectively). Scale bar, 50 μm. **e,** Immunoblot analysis reveals increased tau phosphorylation at S396 in PSP-RS, which is significantly reduced by treatment with LY294002 (PI3K inhibitor) or Torin1 (mTOR inhibitor) (**** P < 0.0001, one-way ANOVA with Tukey’s post hoc test). **f,** Representative images of DA neurons stained for dendritic marker MAP2 (red) and TAU (green) show aberrant dendritic TAU localization in PSP-RS. Scale bar, 25 μm. **g,** Quantification of MAP2⁺ dendrites positive for TAU shows significantly increased dendritic mislocalization in PSP-RS neurons compared to HC (**** P < 0.0001, Mann–Whitney test; n = 19 cells). **h,** LY294002 or Torin1 treatment reduces dendritic TAU accumulation in PSP-RS DA neurons. Scale bar, 25 μm. **i,** Quantification of TAU⁺/MAP2⁺ dendritic segments per cell confirms that both LY294002 (n = 50 PSP cells, 41 LY-treated cells) and Torin1 (n = 28 cells per group) significantly rescue dendritic TAU mislocalization (**** P < 0.0001, Mann–Whitney test). Statistics are available in Additional File 2.

### Pathological matrix remodeling compromises cytoskeletal integrity via RhoA-ROCK signaling

Genes regulating microtubule dynamics, spindle assembly, and G2/M phase transition were downregulated in PSP-RS (**Supplementary Fig. S7a**), suggesting cytoskeletal instability. In parallel, genes encoding integrin-associated adhesion molecules (e.g., *ICAM5, PDGFRL, EDNRA, PDGFRA*) and focal adhesion components (e.g., TLN1, ILK, ITGB4, LAMA2) were upregulated (**Supplementary Fig. S7b-c**), whereas neuronal adhesion markers essential for neurite extension (NCAM1, L1CAM) were suppressed (**Supplementary Fig. S7d**). Kinesins (*KIF2C, KIF20A, KIF11*), spindle regulators (*CKAP2, CENPE*), and actin-binding proteins (PAK1, ATAT1, DCTN1, MAPRE2, STMN2) were also broadly downregulated (**Supplementary Fig. S7e-f**). Proteomic profiling further supported cytoskeletal disorganization, revealing downregulation of actin-binding proteins including gelsolin (GSN), palladin (PALLD), and profilin-2 (PFN2) (**Supplementary Fig. S7g**). Phalloidin staining revealed enhanced F-actin signal in PSP-RS MOs (**Fig. 5a-b**), consistent with filament hyperpolymerization or reduced turnover ^57^. This phenotype was accompanied by reduced expression of CapZ, an actin-capping protein (**Fig. 5c**; **Supplementary Fig. S7h**), and decreased acetylation of α-tubulin at Lys40 (**Fig. 5d; Supplementary Fig. S7i**), indicative of destabilized microtubules ^58^. RhoA–ROCK signaling, a key regulator of actin dynamics ^59–63^, was hyperactivated in PSP-RS organoids, as evidenced by elevated RhoA-GTP levels (**Fig. 5e**). ROCK inhibition with Y-27632 normalized F-actin levels (**Fig. 5f**). Conversely, treatment of HC MOs with recombinant TGF-β isoforms for two weeks induced F-actin accumulation (**Fig. 5g**), whereas blockade of TGF-β receptor signaling with SB43154 reduced actin polymerization in PSP-RS MOs (**Fig. 5h**). Together, these findings establish the TGF-β-RhoA-ROCK axis as a mechanistic link between ECM remodeling and cytoskeletal disorganization ^59–63^.

**Fig. 5.**
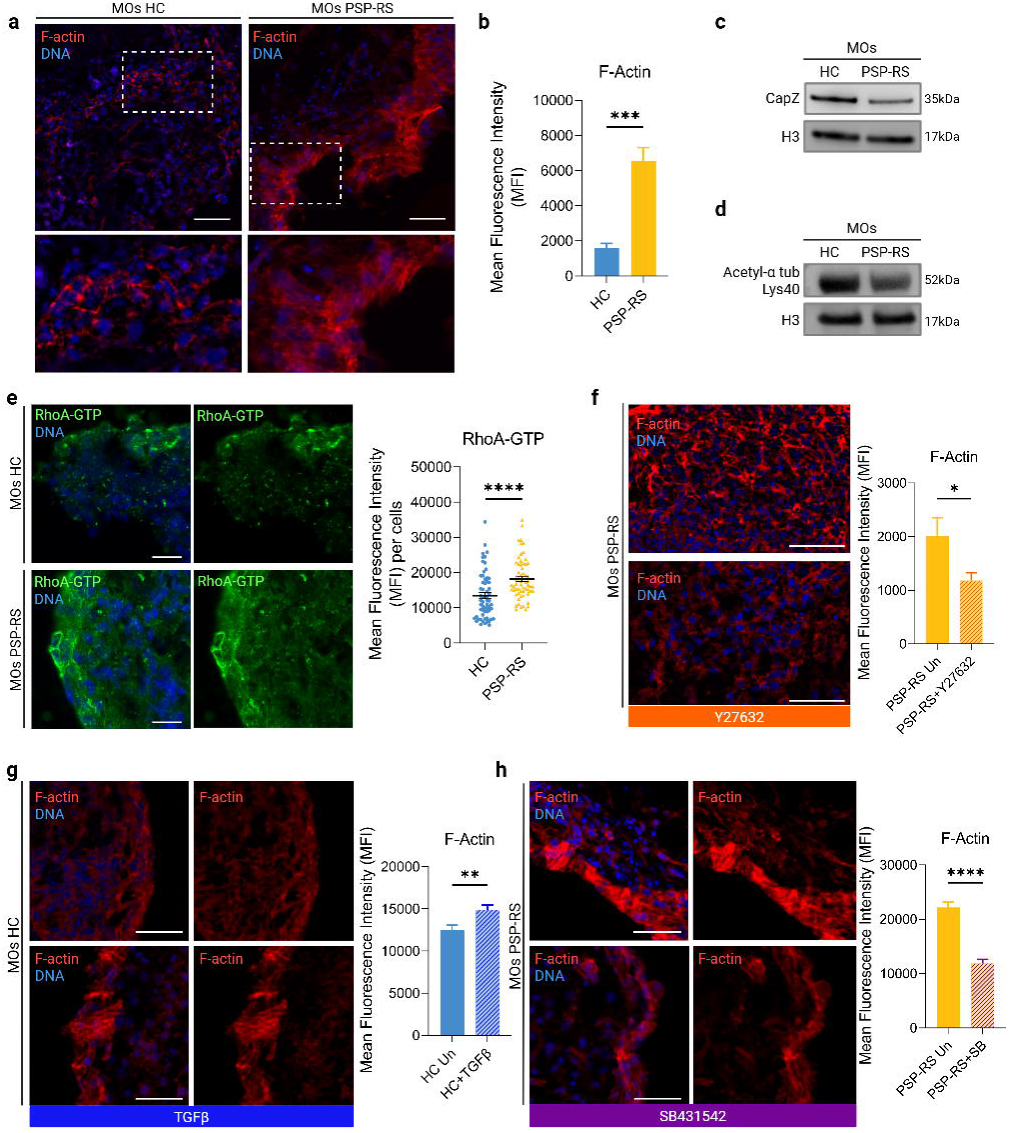
RhoA–ROCK mediates cytoskeletal remodeling downstream of TGF-β–ECM cues. **a,** Representative images of F-actin staining (phalloidin, red) in MOs primary cultures from healthy controls (HC) and PSP-RS patients. DNA is counterstained in blue (DAPI). Insets show higher magnification of boxed areas. Scale bar, 50 µm. **b,** Quantification of mean fluorescence intensity (MFI) for F-actin reveals significantly increased actin polymerization in PSP-RS (mean ± s.e.m.; *n* = 8 images; *** *P* = 0.0002, unpaired *t*-test with Welch’s correction). **c,** Western blot showing reduced expression of actin-capping protein CapZ in PSP-RS compared to HC. **d,** Western blot of acetylated α-tubulin (Lys40) shows decreased levels in PSP-RS MOs. Uncropped western blot gels are available in Source Data **e,** Representative images and quantification of RhoA-GTP levels (green) show increased active RhoA in PSP-RS cells (mean ± s.e.m.; *n* = 60 cells HC, 61 cells PSP; **** *P* < 0.0001, Mann–Whitney test). DNA, DAPI (blue). Scale bar, 25 µm. **f,** Treatment of PSP-RS MOs with ROCK inhibitor Y-27632 reduces F-actin intensity (mean ± s.e.m.; *n* = 8 images HC, 7 images PSP; *P* = 0.0490, unpaired *t*-test with Welch’s correction). Scale bar, 50 µm. **g,** Exogenous TGF-β treatment of HC MOs increases F-actin intensity (mean ± s.e.m.; *n* = 23 ROIs HC, 24 ROIs HC+TGFβ; ** *P* = 0.0052, unpaired *t*-test with Welch’s correction). Scale bar, 50 µm. **h,** TGF-β I receptor inhibitor SB431542 reduces F-actin intensity in PSP-RS cultures (mean ± s.e.m.; *n* = 23 ROIs PSP, 26 ROIs PSP+SB; **** *P* < 0.0001, unpaired *t*-test with Welch’s correction). Scale bar, 50 µm. Statistics are available in Additional File 2.

### ECM-kinase signaling drives synaptic loss and dendritic degeneration

Proteomics profiling revealed downregulation of pathways involved in synapse assembly, vesicle trafficking, axon development, and plasticity including RER1, NFASC, SLIT1, SYT2, SLC17A6, MAPRE2, SYNGR1, SYP, and SYT4 (**Supplementary Fig. S8a-e**). These findings were corroborated by decreased expression of synaptophysin (SYP) (**Supplementary Fig. S8f)**. Immunofluorescence revealed reduced SYP and PSD95 puncta in PSP-RS organoids (**Fig. 6a-b**), consistent with loss of pre- and postsynaptic structure ^24^, along with reduced drebrin, an F-actin– binding protein enriched in dendritic spines (**Fig. 6c**). Morphometric analysis of MAP2-positive neurons revealed shortened neurites and reduced branching complexity (**Fig. 6d-e**), indicative of dendritic atrophy. Strikingly, TGF-β receptor inhibition restored dendritic complexity (**Fig. 6f-g**), whereas exposure of HC MOs to recombinant TGF-β1 recapitulated synaptic and dendritic deficits (**Fig. 6h-i**) seen in disease MOs. Thus, TGF-β–driven ECM remodeling undermines cytoskeletal and synaptic integrity, completing the cascade from VLMC expansion to structural degeneration.

**Fig. 6.**
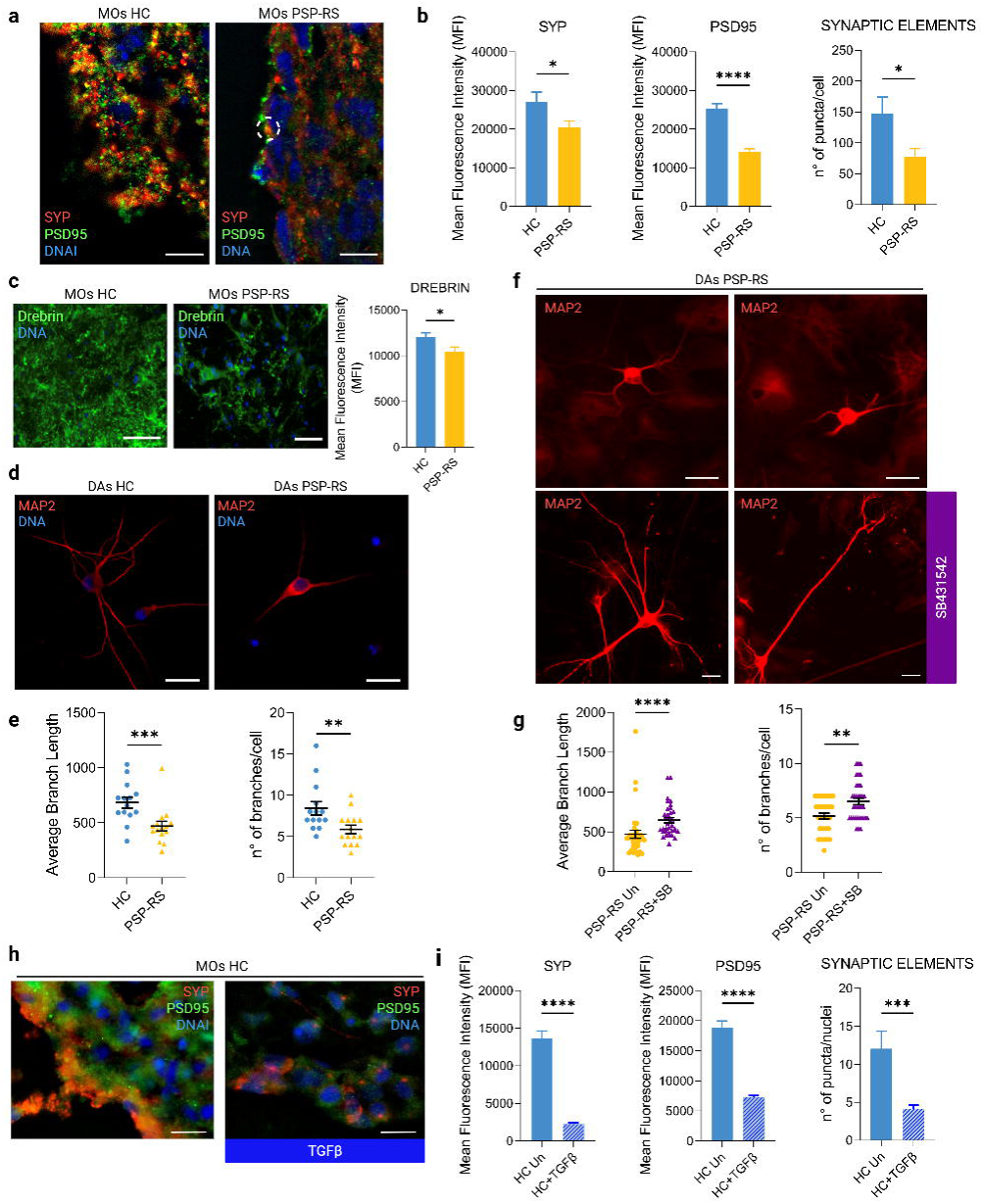
TGF-β–ECM signaling disrupts synaptic architecture and dendritic complexity. **a,** Representative confocal images of MOs from HC and PSP-RS donors stained for synaptophysin (SYP, red), PSD95 (green), and DNA (DAPI, blue) show reduced pre- and post-synaptic puncta in PSP-RS. Scale bar, 10 μm. **b,** Quantification of synaptic markers reveals decreased fluorescence intensity for SYP (P = 0.0273) and PSD95 (**** P < 0.0001), along with fewer synaptic elements per nucleus (P = 0.0362) in PSP-RS (mean ± s.e.m.). **c,** Expression of the dendritic spine marker drebrin is significantly reduced in PSP-RS MOs (mean ± s.e.m.; n = 17 images; P = 0.0210). Scale bar, 50 μm. **d,** Representative images of DA neurons stained for MAP2 (red) reveal diminished neurite complexity in PSP-RS cultures. Scale bar, 25 μm. **e,** Quantification shows significantly shorter dendritic branch length (*** P = 0.0007) and fewer branches per neuron (** P = 0.0074) in PSP-RS DA neurons (mean ± s.e.m.; n = 14–15 neurons). **f, g,** Treatment of PSP-RS neurons with the TGF-β receptor inhibitor SB431542 rescues dendritic complexity, increasing both branch length (**** P < 0.0001) and number of branches per neuron (*** P = 0.0005; n = 35 neurons). Scale bar, 25 μm. **h,** Exogenous TGF-β treatment of HC MOs reduces SYP and PSD95 puncta, mimicking the PSP-RS synaptic phenotype. Scale bar, 25 μm. **i,** Quantification confirms significant reductions in SYP (**** P < 0.0001), PSD95 (**** P < 0.0001), and synaptic elements per nucleus (*** P = 0.0005) following TGF-β treatment (mean ± s.e.m.; n = 30–32 regions of interest). Statistics are available in Additional File 2.

## Discussion

Our findings identify aberrant aberrant TGF-β-driven ECM remodeling as an upstream and sustaining trigger of neurodegeneration in sporadic PSP-Richardson syndrome (PSP-RS). This mechanism reframes disease initiation as a multicellular process in which non-neuronal vascular niche cells, rather than neurons alone, orchestrate pathogenic tau accumulation. Notably, our prior work demonstrated elevated active TGF-β levels in neuronal-derived extracellular vesicle from PSP-RS patients ^18^, further supporting a mechanistic role. By integrating multi-omics with functional assays in patient-derived midbrain organoids, we delineate a pathological cascade linking extracellular structural changes to intracellular signaling, cytoskeletal destabilization, proteostatic failure, and synaptic degeneration. A central insight is that ECM restructuring in not a secondary response to neuronal injury but a primary driver that sustains degeneration through a feedforward circuit of TGF-β activation and integrin clustering. This altered matrix microenvironment engages RhoA-ROCK-mediated cytoskeletal remodeling ^59,61^, and sustains PI3K/AKT and MAPK/ERK signaling. Both cascades drive tau hyperphosphorylation through kinase activation and PP2A suppression ^49,50^. The parallel engagement of AKT and ERK indicates convergent, rather than redundant, inputs into tau pathology, suggesting that their simultaneous inhibition may provide superior therapeutic benefit. Downstream of AKT, we identify mTORC1 hyperactivation as a proteostatic checkpoint linking ECM signaling to autophagy impairment. This dysregulation drives p62/SQSTM1 accumulation and tau buildup ^64^, a phenotype sufficient to induce tau accumulation even in otherwise healthy neurons ^65^. Pharmacological mTORC1 inhibition restores autophagic flux and reduces tau burden, supporting a model in which impaired degradation, rather than enhanced production, is the dominant driver of tauopathy ^56^. Importantly, these intracellular defects culminate in structural synaptic compromise, marked by loss of pre- and postsynaptic markers, dendritic atrophy, and reduced branching complexity. Pharmacological blockade of TGF-β signaling restored both autophagy and neuronal architecture, underscoring the causal role of ECM remodeling in structural degeneration ^66^. Single-cell RNA sequencing further anchored these findings by revealing expansion of collagen-secreting vascular leptomeningeal-like cells (VLMCs) and transcriptional reprogramming across non-neuronal lineages, including stress responses in progenitors and astrocytes ^41^. This broad multicellular response suggests that VLMC-derived matrix alterations act as upstream cues that condition the neural microenvironment toward degeneration. Collectively, our results position the non-neuronal TGF-β-matrix axis as a pathogenic convergence point integrating structural, transcriptional, and proteostatic dysfunction in PSP-RS. By highlighting VLMC-mediated ECM remodeling as a generalizable initiating mechanism, this work reframes vascular riche cells as active drivers of neurodegeneration. This framework extends beyond PSP-RS, as ECM-TGF-β signatures are detectable in other tauopathies, offering multiple therapeutic entry points to intercept tau pathology at an early, potentially reversible stage before irreversible neuronal loss occurs.

## Acknowledgements

We are deeply grateful to the donors whose generous contributions made this work possible. We thank the Proteomic Facility of the University of Catanzaro led by M.G. (proteomicsumg.org) for technical support, and the Novogene Core Facility for assistance with bulk RNA transcriptomic experiments. Single-cell RNA-sequencing (10x Genomics) was performed by Negedia S.r.l. We also thank Anna Maria Aloisio for technical assistance. Illustrations were created with BioRender.com.

## Author contributions

E.I.P. and G.C. conceptualized and supervised the overall project. C.Z., D.V., D.B., A.B., S.S., C.G., R.C., G.L.B., M.T., V.A., A.F., performed experiments. C.Z., D.V., D.B., A.B., An.Q., V.A., and S.S. analyzed data. C.Z., D.V., D.B, A.B., and E.I.P. prepared figures. G.C. and E.I.P wrote the manuscript with assistance from A.F, F.C., Al.Q, and An.Q and input from all authors. D.B. and F.C. performed bioinformatic analysis. C.G. and M.G. carried out mass spectrometry. Al.Q. and An.Q. recruited the PSP-RS patients and healthy subjects used in this study.

## Data availability

This work did not produce original code. All data supporting the findings of this work are available within the main text, Extended Data Figures, and Supplementary Information. Bulk RNA and single cell RNA-seq data, mass spectrometry proteomics datasets, and whole-genome sequencing data have been deposited in public repositories ^67, 68^ and will be made accessible upon publication. Source data underlying all bar plots, as well as uncropped scans of immunoblots and gels, are provided. Unique biological materials generated in this study are available from the corresponding author upon reasonable request without restrictions.

## Declarations

## Ethics approval and consent to participate

Study protocols, amendments, and informed consent forms were reviewed and approved by local institutional review boards/independent ethics committees. Written informed consent was obtained from each participant.

## Consent for publication

Not applicable.

