## Supplementary figures and images for "Non-neuronal, TGF-β–driven extracellular matrix restructuring promotes neurodegeneration in a PSP-Richardson syndrome model"

### Fig S1.png

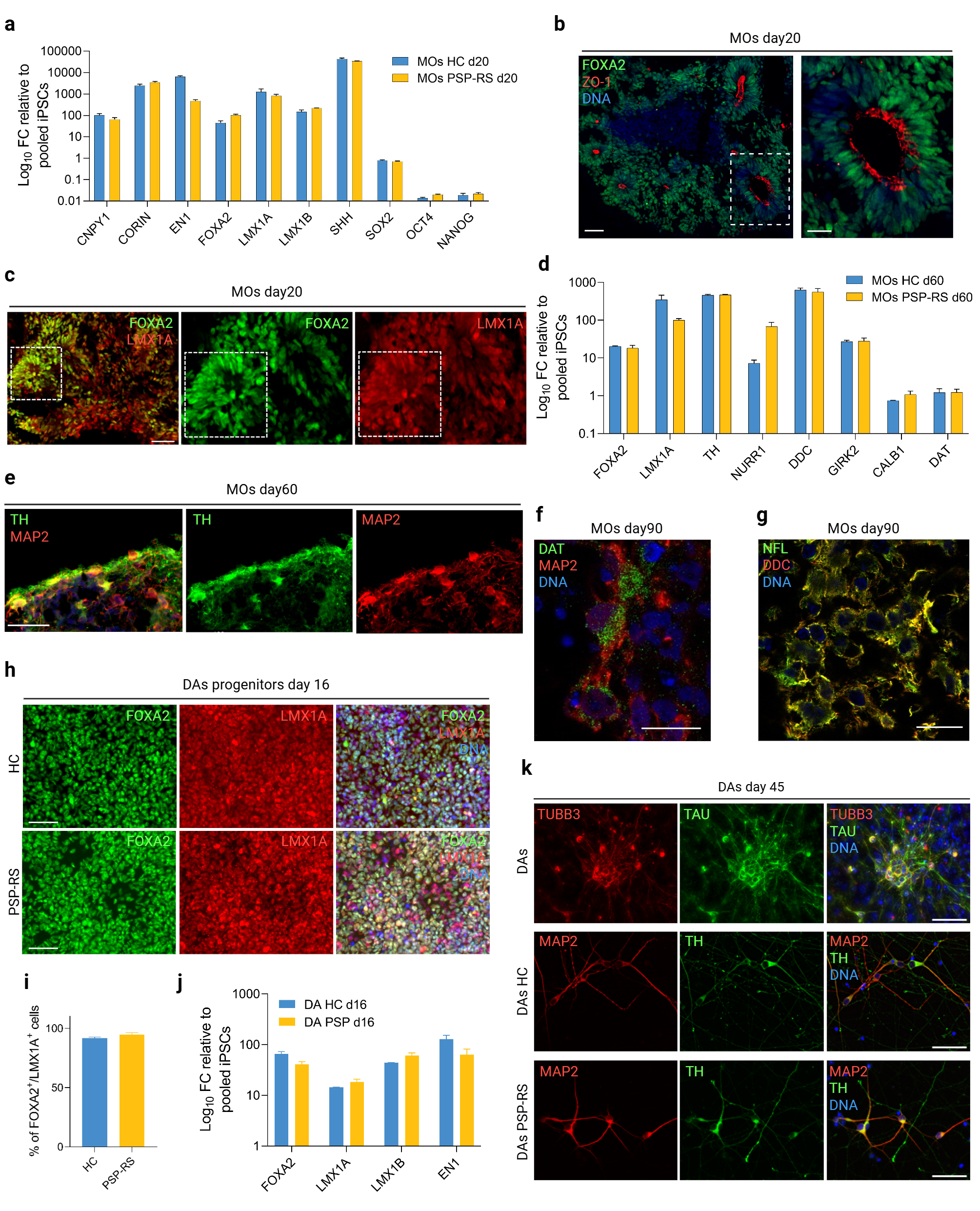

### Fig S2.png

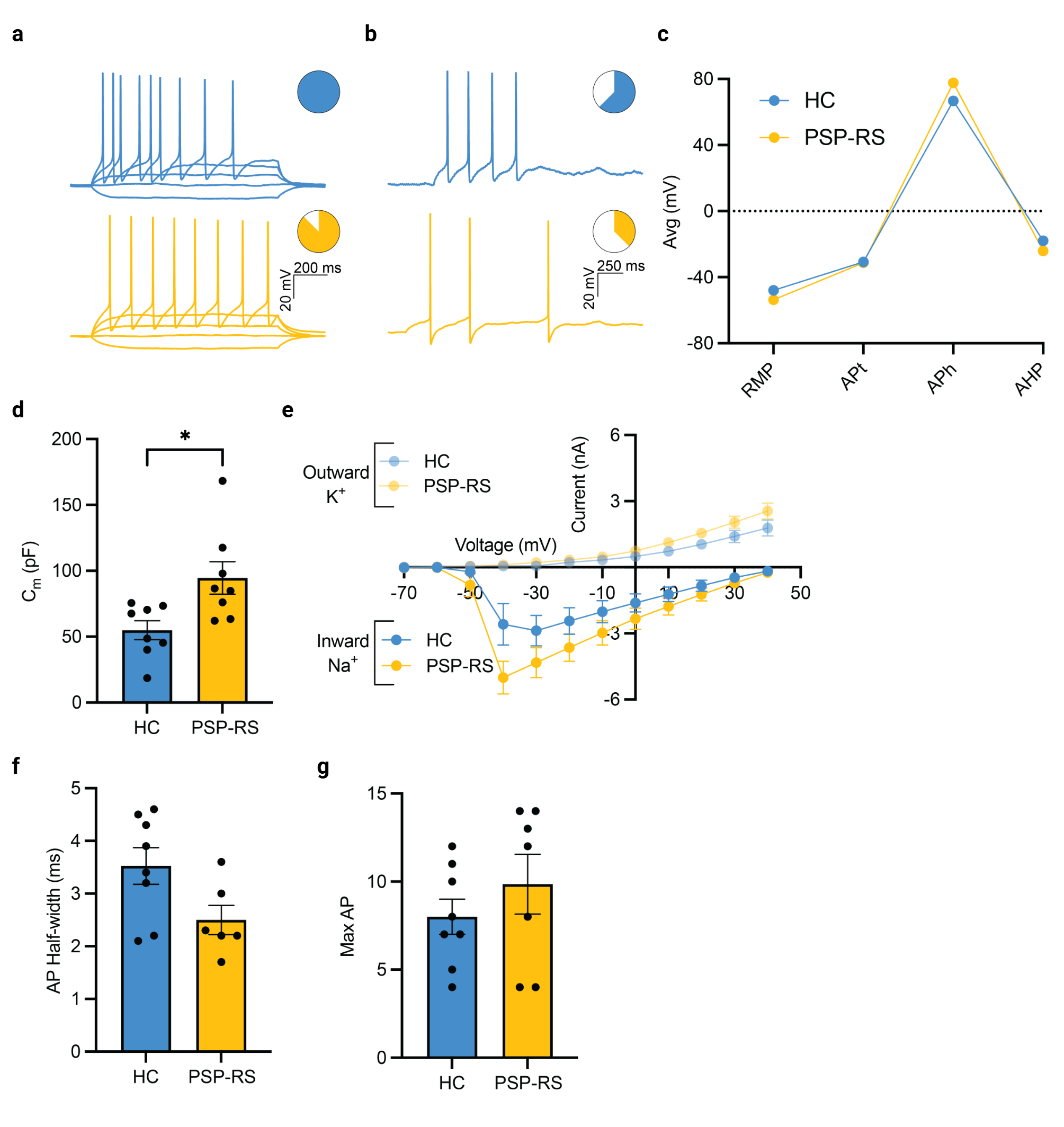

### Fig S3.png

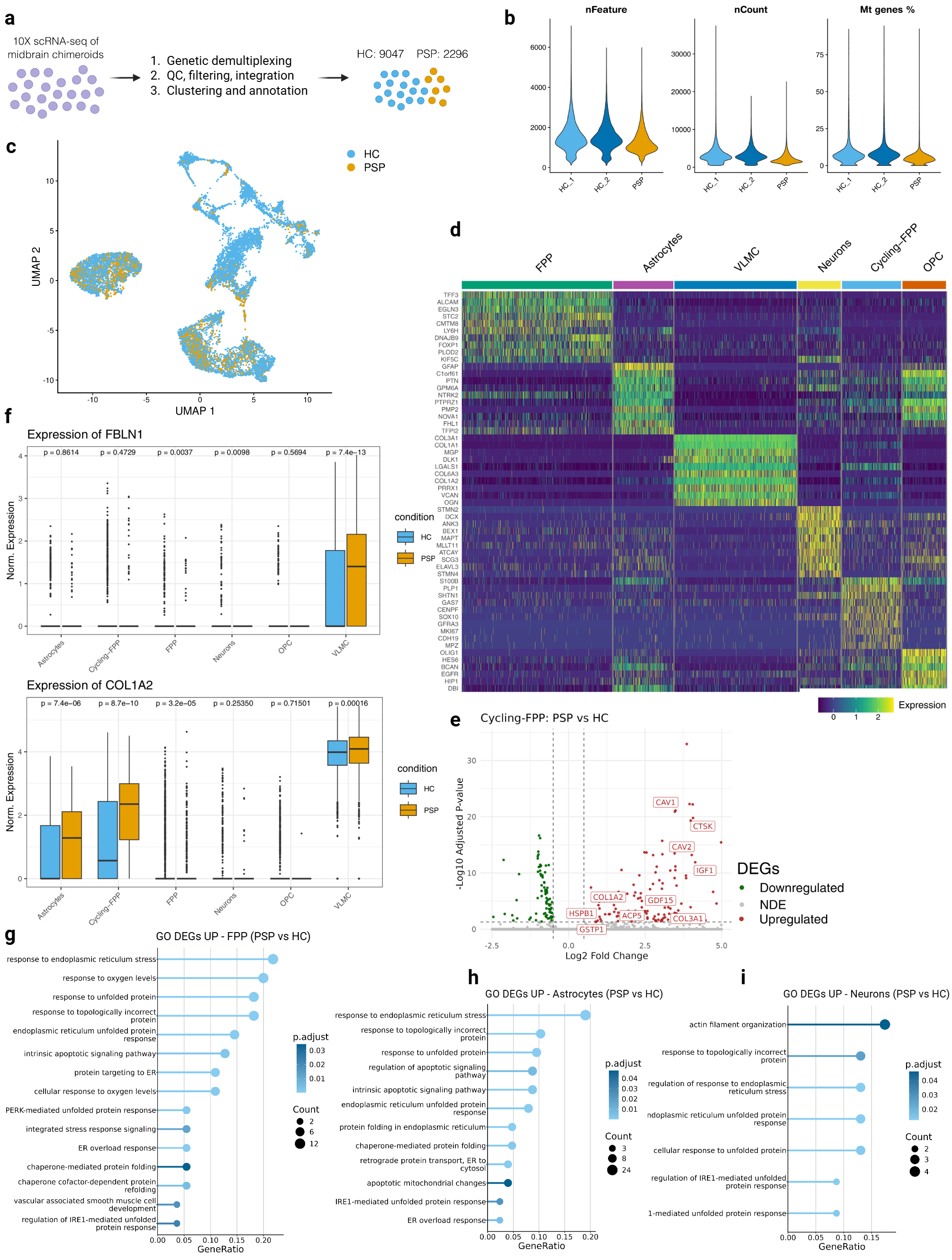

### Fig S4.png

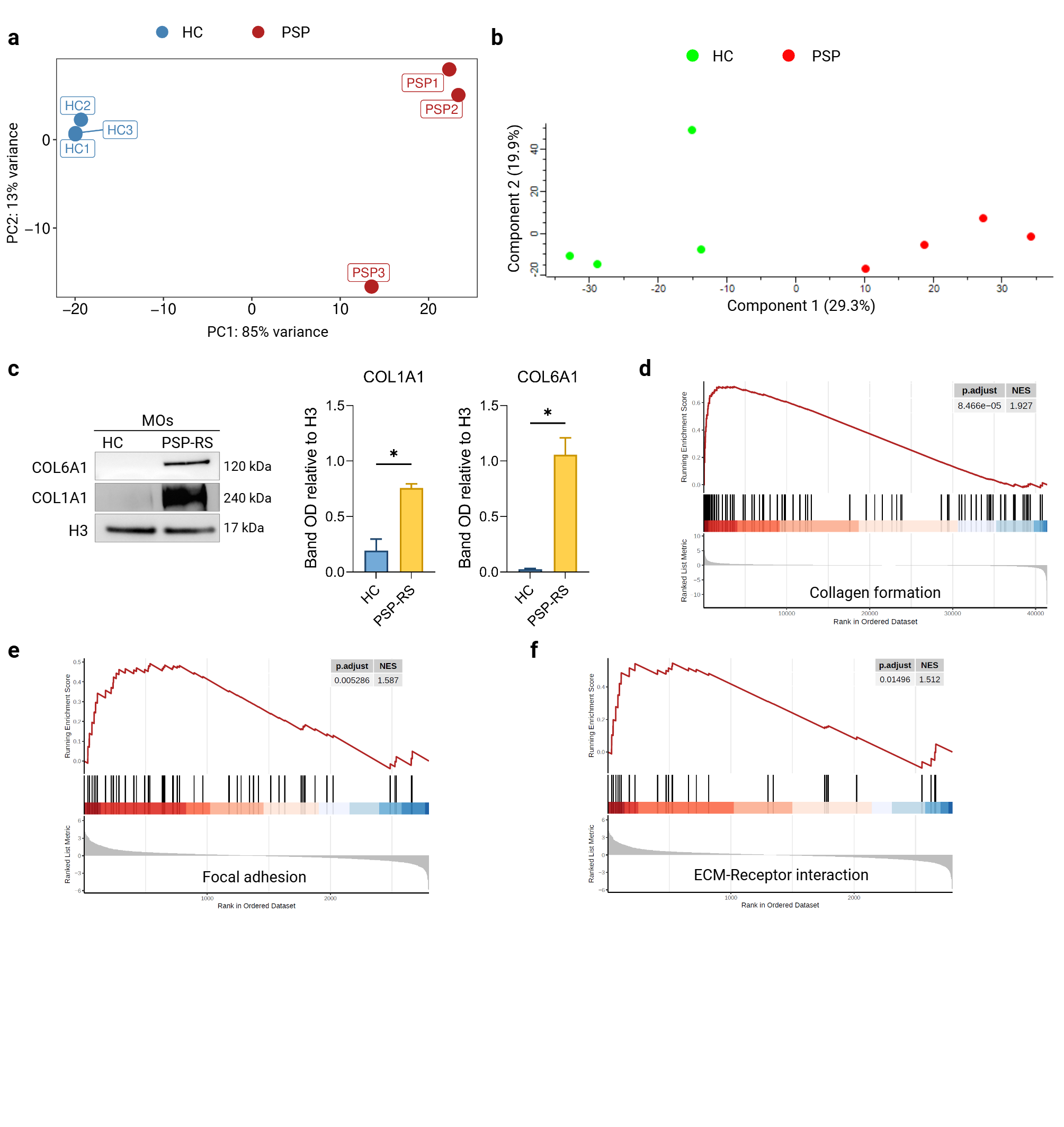

### Fig S5.png

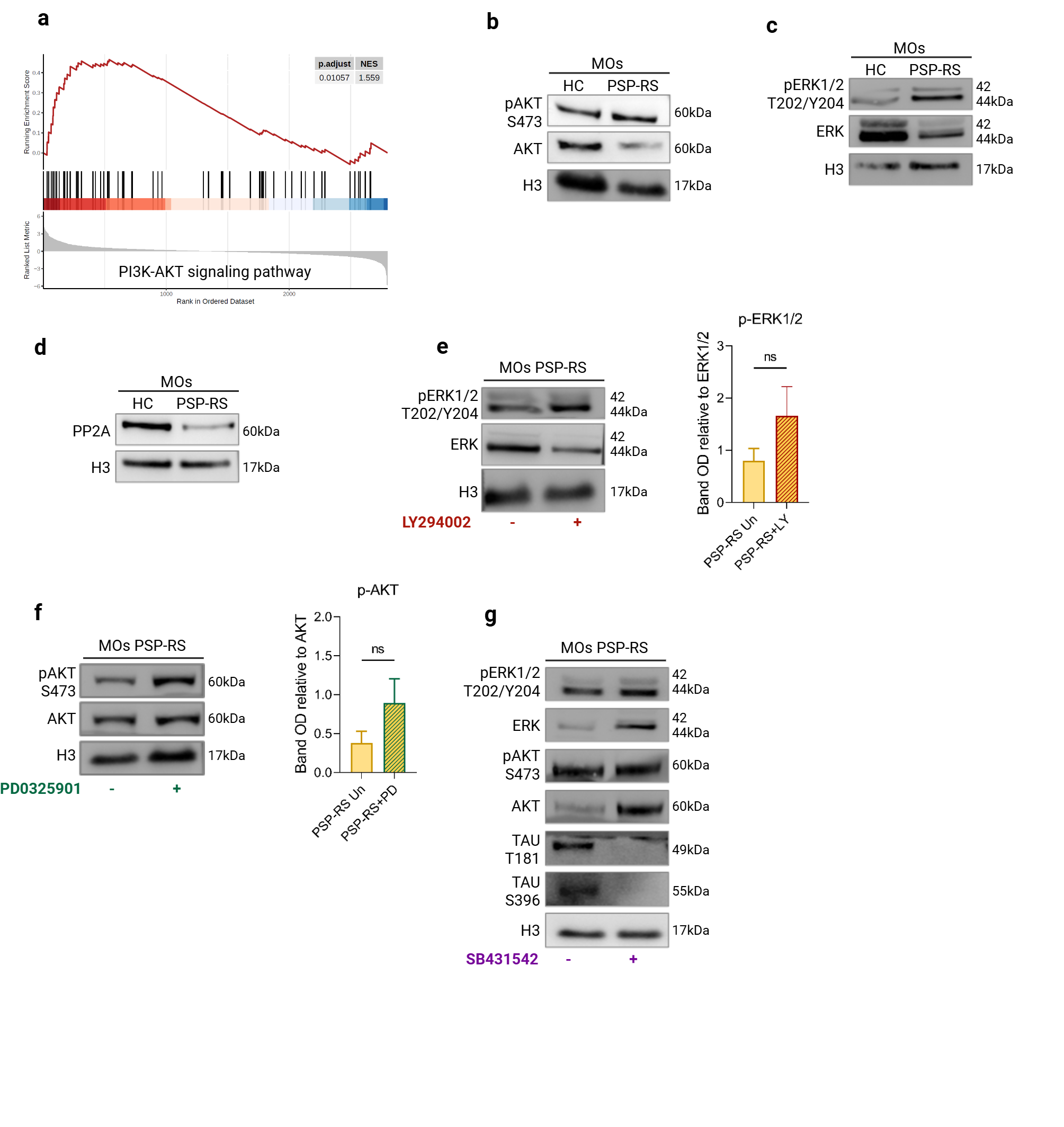

### Fig S6.png

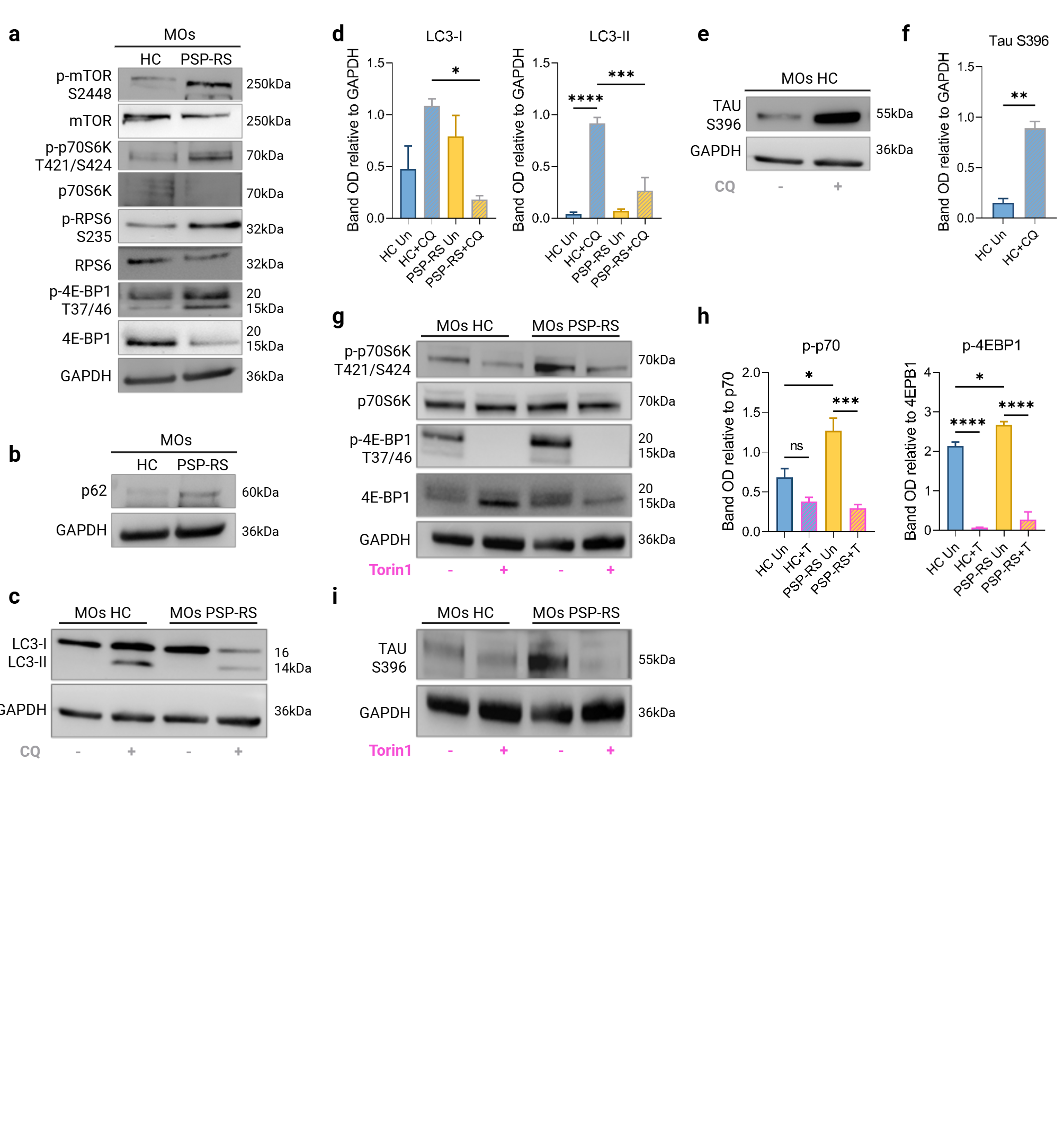

### Fig S7.png

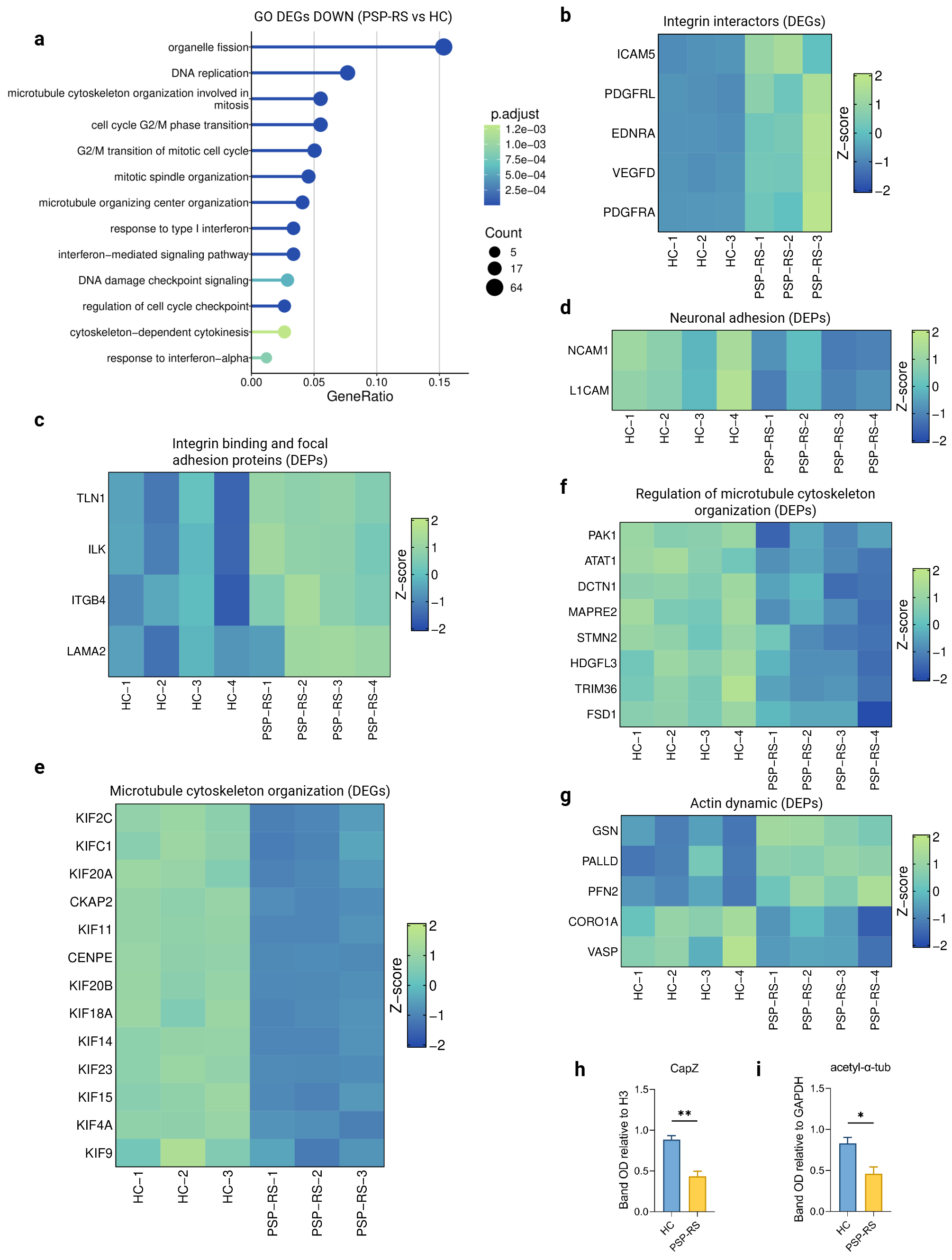

### Fig S8.png

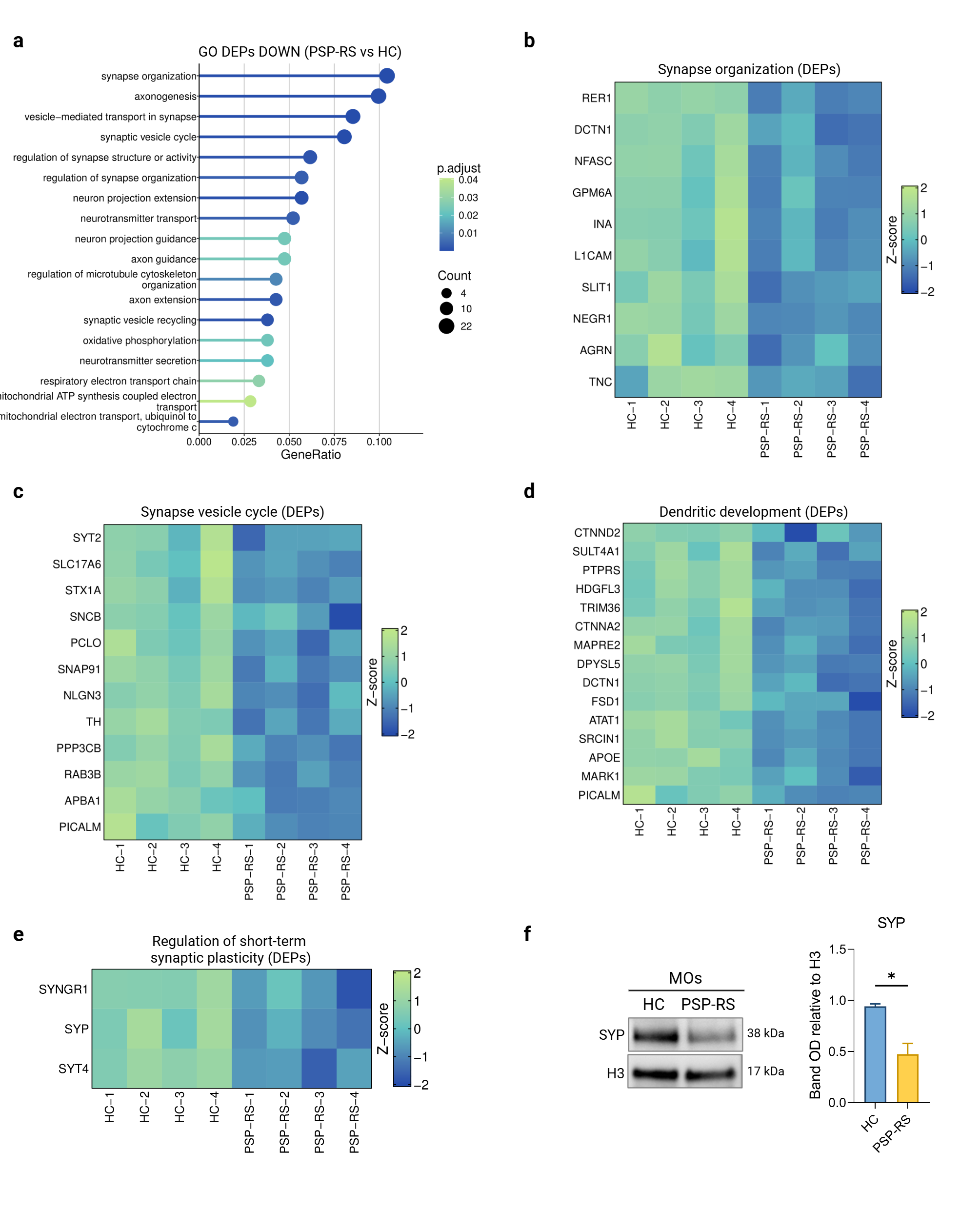

### Supplementary Data S1.png

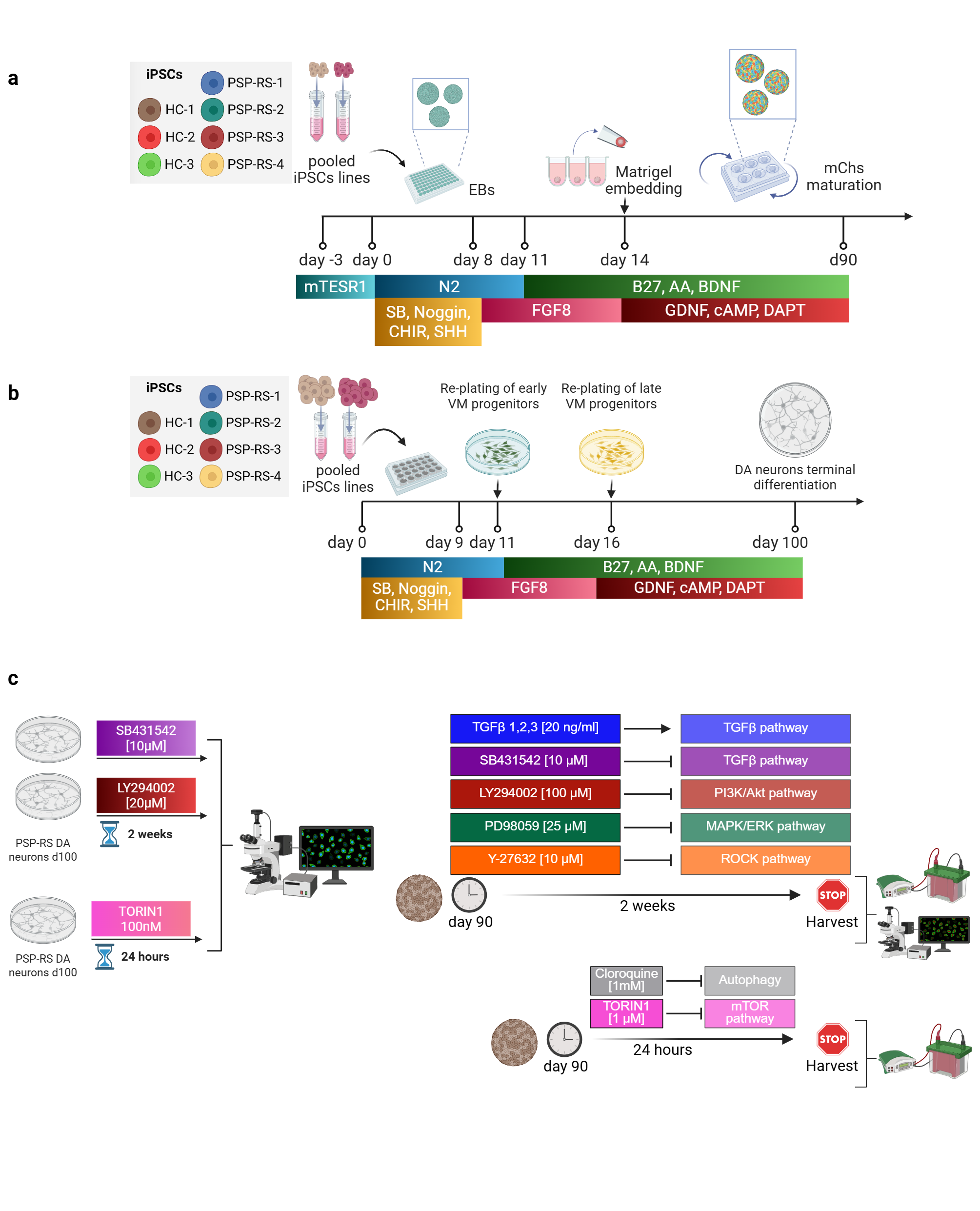
